# Multi-level force-dependent allosteric enhancement of αE-catenin binding to F-actin by vinculin

**DOI:** 10.1101/2022.05.07.491039

**Authors:** Nicolas A. Bax, Amy Wang, Derek L. Huang, Sabine Pokutta, William I. Weis, Alexander R. Dunn

**Author notes:** These authors contributed equally to this study (Nicolas A. Bax, Amy Wang).

## Abstract

Classical cadherins are transmembrane proteins whose extracellular domains link neighboring cells, and whose intracellular domains connect to the actin cytoskeleton via β-catenin, α- catenin. The cadherin-catenin complex transmits forces that drive tissue morphogenesis and wound healing. In addition, tension-dependent changes in αE-catenin conformation enables it to recruit the actin-binding protein vinculin to cell-cell junctions, where it contributes to junctional strengthening. How and whether multiple cadherin-complexes cooperate to reinforce cell-cell junctions in response to load remains poorly understood. Here, we used single-molecule optical trap measurements to examine how multiple cadherin-catenin complexes interact with F-actin under load, and how this interaction is influenced by the presence of vinculin. We show that force oriented toward the (-) end of the actin filament results in mean lifetimes 3-fold longer than when force was applied towards the barbed (+) end. Further, load is distributed asymmetrically among complexes, such that only one bears the majority of applied load. We also measured force-dependent actin binding by a quaternary complex comprising the cadherin-catenin complex and the vinculin head region, which cannot itself bind actin. Binding lifetimes of this quaternary complex increased as additional complexes bound F-actin, but only when load was oriented toward the (-) end. In contrast, the cadherin-catenin complex alone did not show this form of cooperativity. These findings reveal multi-level, force-dependent regulation that enhances the strength of the association of multiple cadherin/catenin complexes with F-actin, conferring positive feedback that may strengthen the junction and polarize F-actin to facilitate the emergence of higher-order cytoskeletal organization.

## Introduction

Classical cadherins are transmembrane proteins that mediate homophilic interactions between cells, and are fundamental to the construction of animal tissues [1, 2]. Cadherins are linked to the underlying actomyosin cytoskeleton by β-catenin, which binds to the cadherin cytoplasmic tail and to αE-catenin, which in turn binds to F-actin [3, 4] (Fig. 1A). This molecular linkage maintains tension at cell-cell contacts [5, 6], and is essential for dynamic mechanical coupling between cells during morphogenesis and for tissue homeostasis [1, 5, 7–10]. During these and other processes, cell-generated forces must be coordinated and transmitted across tissues. In particular, cables of contractile filamentous (F)-actin and nonmuscle myosin II spanning multiple cells drive large-scale tissue rearrangements during embryonic development and wound healing [11–15]. The sarcomeric actomyosin arrays that power muscle contraction in the heart are similarly linked by cadherin-catenin complexes at cardiomyocyte cell-cell junctions. How these intercellular connections self-assemble, and how they remain stable under mechanical load, is unclear.

**Fig. 1.**
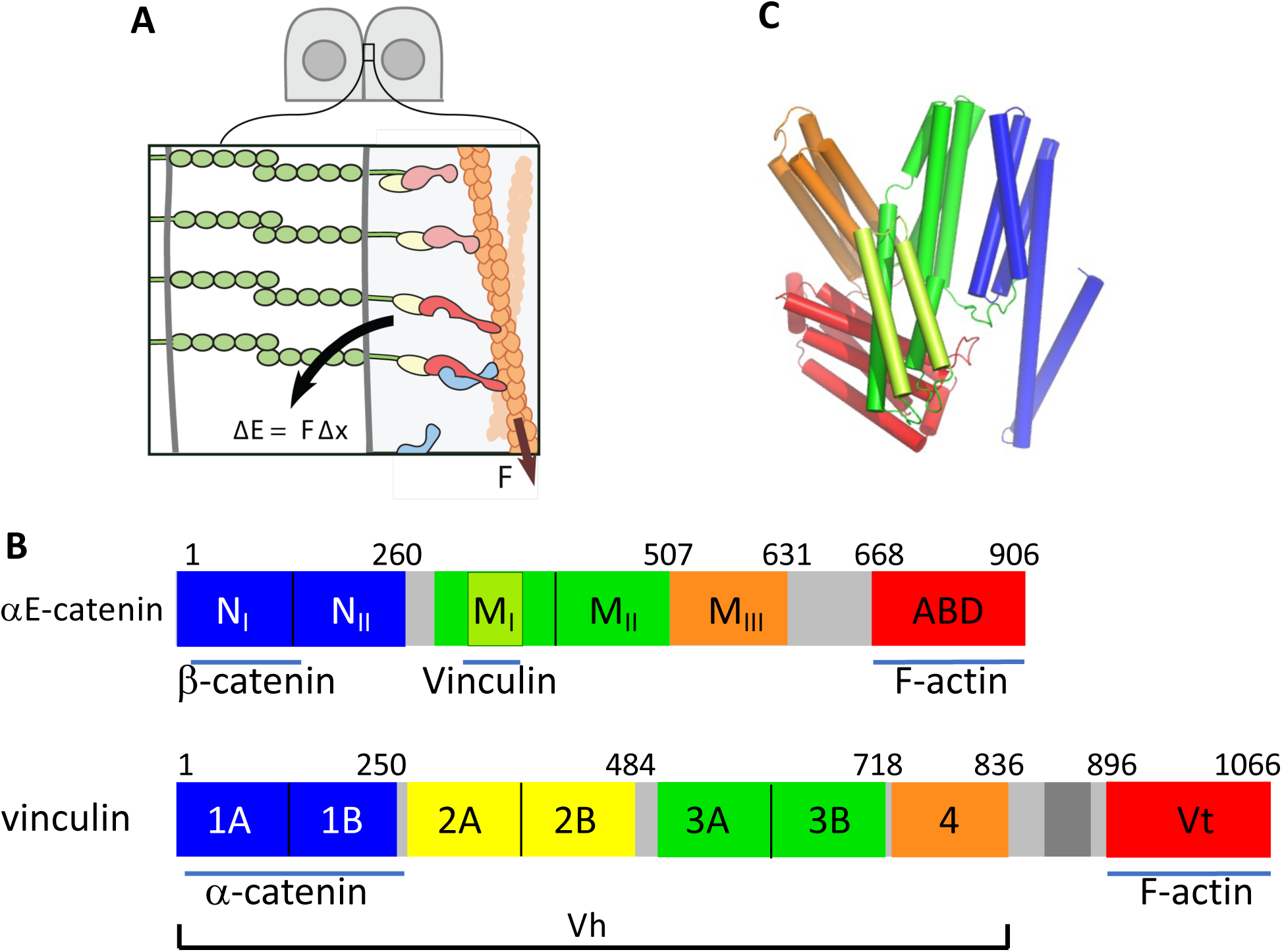
αE-catenin and vinculin at cell-cell contacts. **(A)** Schematic of a minimal cell-cell contact containing a classical cadherin (green), β-catenin (yellow), αE-catenin (red) and actin. Vinculin (blue) is recruited to the contact upon application of tension to αE-catenin. (**B)** Primary structures of αE-catenin (*top)* and vinculin *(bottom).* The N-terminal domain of αE-catenin, which binds β-catenin, contains two four-helix bundles, N_I_ and N_II_ [40, 64–66]. The M domain consists of three four-helix bundles, designated M_I_, M_II_ and M_III_. The C-terminal ABD is a five-helix bundle. The vinculin N-terminal D1 domain confers full binding affinity for αE-catenin [26]. The head region spans domains 1-4. **C)** Crystal structure of αE-catenin 82-906 [65].

The ternary epithelial (E)-cadherin/β-catenin/αE-catenin complex binds weakly to F-actin [16, 17], but binding is strengthened by mechanical force, a property known as a catch bond, which is thought to reinforce intercellular contacts under tension [18]. In addition to the minimal ternary cadherin/β-catenin/α-catenin complex, other proteins bind to αE-catenin and F-actin depending on the mechanical environment of the junction [19–21]. The best-studied example is vinculin, a paralog of αE-catenin found in focal adhesions and cell-cell junctions (Fig. 1B).

Vinculin is recruited to cell-cell junctions upon application of force to αE-catenin [22–24], where it strengthens the adhesive contact between cells [25]. In solution, both the N-terminal D1 domain of vinculin and the larger vinculin “head” (designated Vh) comprising all but its actin-binding domain (Fig. 1B), bind αE-catenin or its complex with E-cadherin and β-catenin weakly (K_D_ = ∼2 μM) whereas both fragments bind to the isolated M_I_-M_II_ fragment of αE-catenin with high affinity (K_D_ = ∼15 nM) [26]. Mechanical tension on αE-catenin is thought to reversibly displace intramolecular interactions within the M domain [27], exposing the αE-catenin M_I_ subdomain and allowing it to bind strongly to vinculin [26, 28]. However, little is known about the consequences of vinculin binding on the interaction of the cadherin-catenin complex with F-actin.

## Results

### The ternary complex shows asymmetric force-dependent binding to actin

We employed an optical trap (OT) assay [18, 29, 30] to compare the behavior of the ternary E-cadherin/β-catenin/αE-catenin complex with the quaternary complex formed by adding the vinculin head (Fig. 2A). In this assay, a biotinylated actin filament is attached to two streptavidin-coated beads, which are each captured in an optical trap. This actin “dumbbell” is positioned over a platform displaying immobilized recombinant cadherin/catenin/(vinculin) complexes (Fig. 2A). The stage is then moved back-and-forth approximately parallel to the filament. If an αE-catenin molecule attaches to the filament, stage motion pulls a bead out of its trap. When a bead displacement is detected, designated here as an “event”, the stage movement is halted, leaving the complex under tension due to the restoring force of the trap, which acts as a simple spring that pulls the bead back to the waist of the laser beam. Note that the assay does not directly detect the presence of binding interactions of “bystander” complexes that bear little or no load, as these would not affect the positions of the optically trapped beads. When a load-bearing complex detaches from the filament, the force from that attachment is lost, thereby providing measures of both the force and how long the attachment lasted. Most events had multiple steps in force before returning to baseline, which correspond to successive releases of individual complexes, such that the last step before all tension is lost corresponds to that of a single load-bearing complex.

**Fig. 2.**
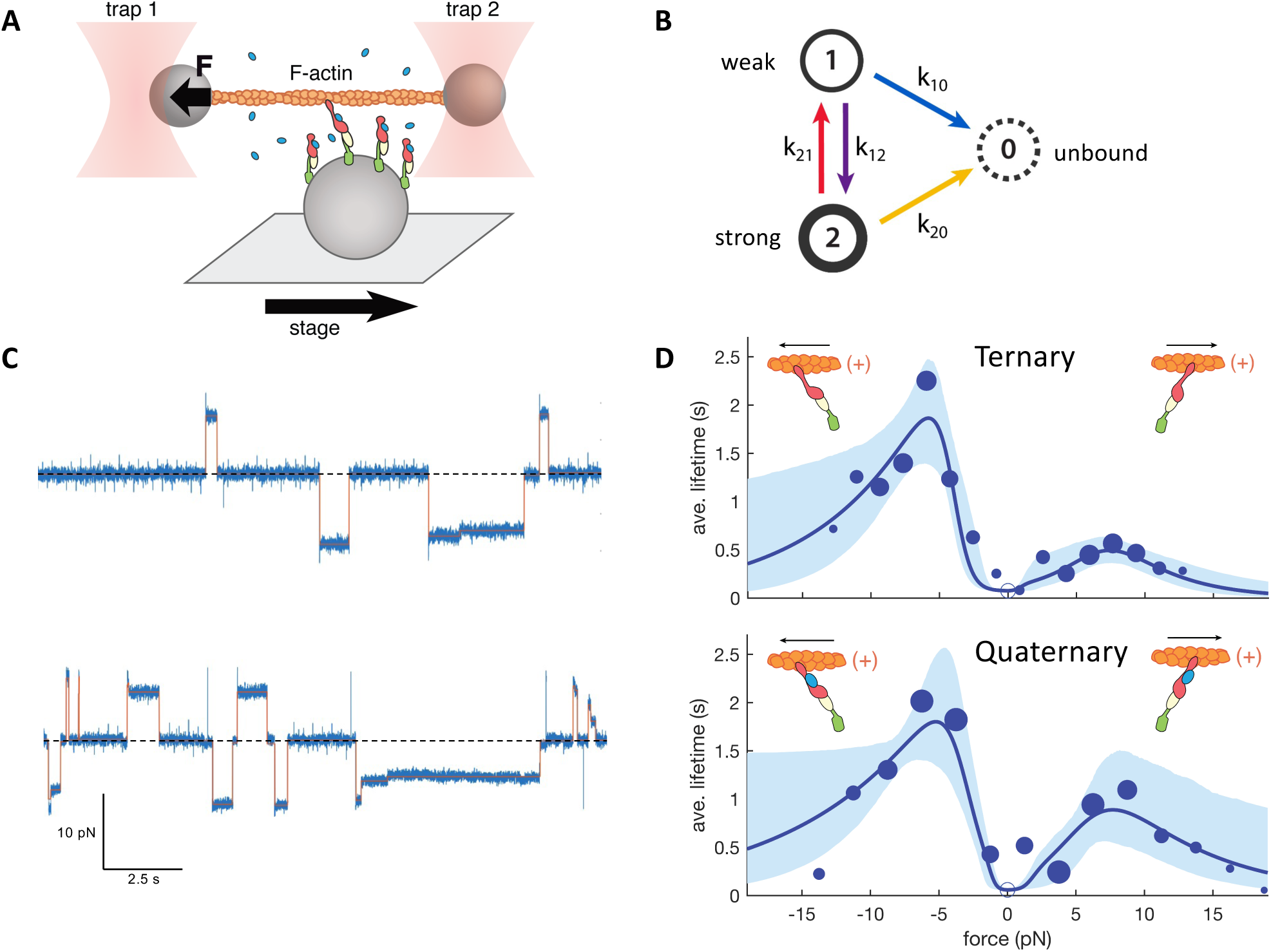
Force-dependent binding of cadherin/catenin complexes to F-actin. **(A)** Schematic of the OT assay. **(B)** The two-state catch bond model used in this work. Weak and strong actin- bound states 1 and 2 can interconvert, and either can dissociate from actin. Force promotes the transition between the weakly bound and strongly bound states, and also disfavors the transition from the strong to the weak state. **(C)** Representative OT data of the ternary (upper) and quaternary (lower) complexes. The zero-force baseline is shown as a dashed line. The assignment of filament direction is described in Supplemental Information. **(D)** Mean binding lifetimes (blue circles) and best-fit model (blue curve) for the ternary E-cadherin cytoplasmic domain/β-catenin/αE-catenin (upper plot) and quaternary E-cadherin cytoplasmic domain/β-catenin/αE-catenin/Vh (lower plot) derived from the last step data. Negative and positive values of force correspond to forces directed towards the (-) or (+) and of the filament, respectively. Areas of the circles are proportional to the number of events measured in each 2 pN bin. Open circle at force = 0 represents the constraint for the binding lifetime at low forces measured using a separate assay. Solid curves are the fit of the two-state catch bond model to the data, and the lighter envelope is the 95% confidence interval obtained by bootstrapping.

The bound lifetimes of single load-bearing complexes had a biexponential distribution at any given force, indicating distinct short- and long-lived states [18, 29]. This observation is consistent with a two-state catch bond [31, 32] in which force increases the rate of formation of a strong-binding state to actin and decreases the back reaction to the weak state (Fig. 2B) [18, 29]. The rates of interconversion between these states are described by the Bell-Evans model [33, 34], wherein the rate of a transition *k_ij_* between states *i* and *j* = *k^0^_ij_* exp (*Fx_ij_* / k_B_T) where *k^0^_ij_* is the rate at zero force, *F* is the force magnitude, and *x_ij_* is projection of the force vector onto the reaction coordinate *r_ij_* (see below). Large values of *x_ij_* indicate a high degree of force sensitivity and imply a large underlying structural transition.

Using an improved OT setup and determining the polarity of the filament by measuring the direction of its movement in a separate flow channel containing the pointed (-) end directed motor myosin VI [29], we demonstrated that vinculin itself shows catch bond behavior that is asymmetric: its lifetime bound to F-actin is longer when force is directed towards the pointed (-) end of the actin filament than toward the barbed (+) end [29]. Asymmetric catch bond behavior was also recently reported for αE-catenin alone [35]. We therefore re-measured the force-dependent association of the ternary complex comprising the E-cadherin cytoplasmic domain, β-catenin, and αE-catenin with F-actin (Fig. 2C). As before, analysis of the last step data revealed biphasic lifetimes at a given force (Fig. S1). The lifetimes of the bound complex increase with force up to about 6-7 pN, and show asymmetry, with longer bound lifetimes when force is directed towards the (-) end (Fig. 2C, D; Supplemental Information). The asymmetric catch bond mechanism was recently rationalized based on the structure of the αE-catenin actin-binding domain (ABD) bound to F-actin [36] and validated in single molecule experiments [30]. We modeled these effects by directionally-dependent distance parameters, *x_ij_^(-)^* and *x_ij_^(+)^*, denoting distance parameters for when force is oriented toward the F-actin (-) or (+) end, respectively (Supplemental Information, Table S1).

### Effect of vinculin on cadherin/catenin complex binding to actin

To assess the effect of vinculin on the actin-binding activity of the ternary cadherin/catenin complex, we performed the OT assay on the quaternary E-cadherin/β- catenin/αE-catenin/vinculin complex made with the vinculin head (Vh), which lacks the vinculin actin-binding domain but contains the binding site for αE-catenin [26] (Fig. 1B, Fig. S2). Vh was added at 15 μM to ensure that the cadherin/catenin complex will be nearly saturated with the vinculin head (see Methods). Analysis of the last step data showed that the vinculin head increased the lifetimes of a single bound complex on F-actin at low forces when load was oriented in the (-) direction, and over a broad range of forces when load was oriented in the (+) direction (Fig. 2C,D, Fig. S3). However, the difference in lifetimes with the ternary complex is relatively modest, and we cannot rule out that an unknown source of systematic error gave rise to this effect.

In cells, the cadherin-catenin complex assembles into large, hierarchically organized clusters [37–39], but how multiple cadherin-catenin complexes might interact when binding to the same actin filament under load has not been examined. For both the ternary and quaternary complexes, when force was directed towards the (+) end of F-actin, the mean lifetime of a complex stayed constant, regardless of how many complexes were bound (Fig. 3A). For the ternary complex, when force was directed towards the (-) end of F-actin, the lifetimes stayed constant or decreased as more complexes were bound. (Fig. 3A; Table S2). In contrast, as more quaternary complexes were bound, the mean binding lifetime of a single complex increased (Fig. 3A; Table S2). The mean forces for each step were similar for both the ternary and quaternary complex (Table S2) and could not explain this observation.

**Fig. 3.**
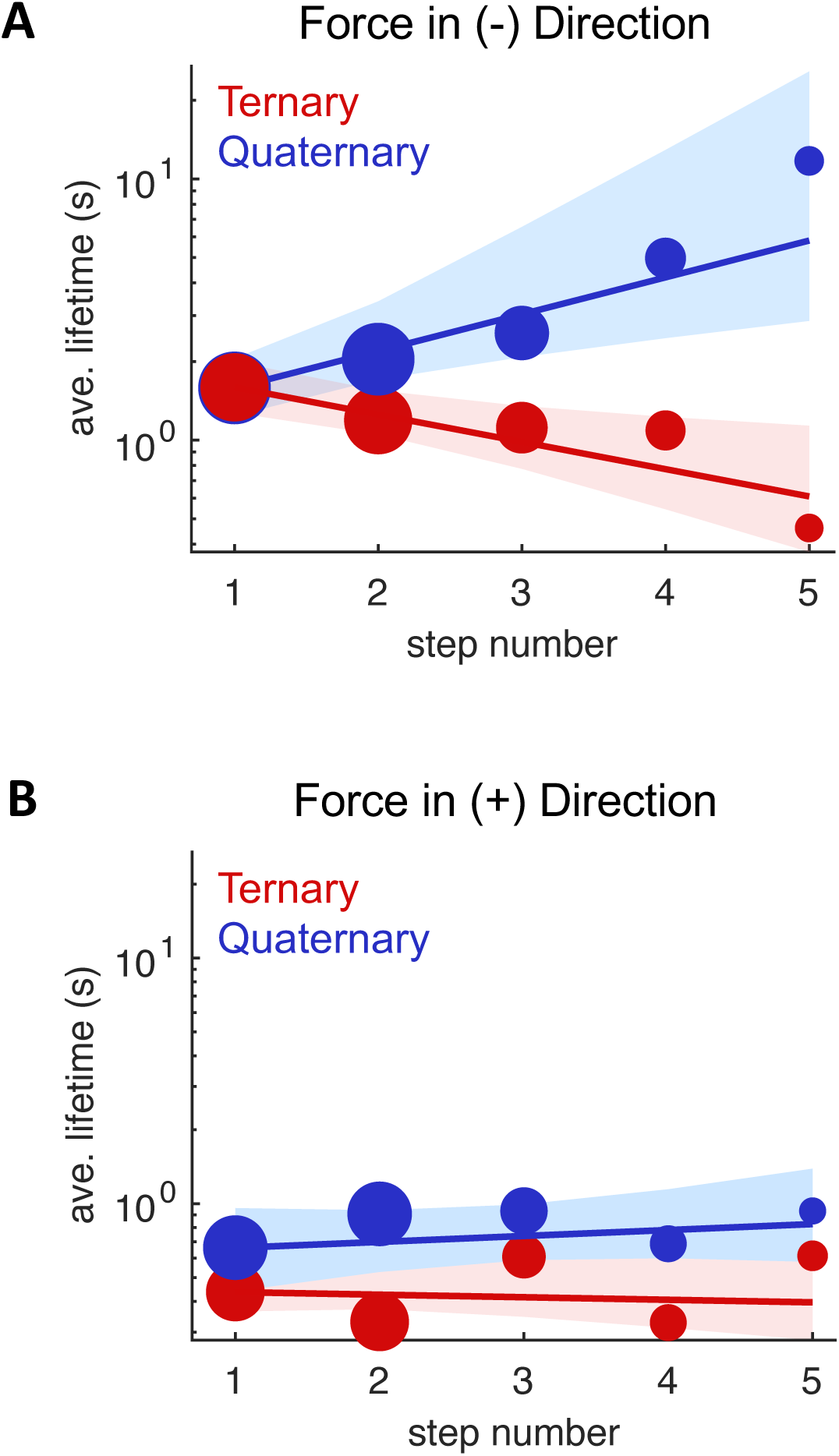
Binding of multiple complexes alters lifetime of individual bonds. **(A, B)** The mean bound lifetime of individual complexes is shown as a function of the number of complexes bound during an event, with load in the (-) and (+) directions, respectively. The size of the circles corresponds to the number of observations. The lines are fit to the function *L*(*n*) = *L*_1_ *exp*(*c* (*n* − 1)., where *L*(*n*) is the expected lifetime for a step number *n*, *L*_1_ is the lifetime of the first-from-end step (*i.e.*, last step), and *c* a constant (see SI text). The envelopes represent the 95% confidence interval obtained by bootstrapping.

Using an assay to measure lifetimes at low forces [29], we found that the vinculin head had little effect on F-actin binding in this regime (SI text and Table S3). We additionally compared the actin-binding activities of the ternary and quaternary complexes in solution to independently assess whether vinculin has any effect on actin binding in the complete absence of load. For these experiments we used the αE-catenin mutant R551A, which disrupts a salt bridge in the M domain and thereby relieves the inhibitory activity of M_III_ toward vinculin binding [40], and vinculin D1, a construct comprising the N-terminal 1A and 1B domains (Fig. 1B), which binds to wild-type αE-catenin with an affinity comparable to that of Vh [26]. The cadherin-catenin complex made with αE-catenin R551A binds strongly to vinculin D1 (K_D_ = 26 nM vs. 1.9 μM for the wild-type ternary complex [41]), ensuring that the complex could be completely saturated in these experiments. Vinculin D1, had no detectable effect on the affinity of the cadherin-catenin complex for F-actin in solution (Fig. S4), confirming that the enhanced lifetimes of the quaternary complex seen in the OT assay are a consequence of applied load. Thus, force not only enables binding of vinculin to αE-catenin, but is also required for vinculin to alter the actin-binding activity of the cadherin/β-catenin/αE-catenin complex.

### Behavior of multiple actin-bound complexes

Structural and biochemical studies [36, 42] found direct contacts between actin-bound αE- catenin ABDs, and suggested that these interactions enhance binding between αE-catenin and F-actin. Cooperative binding of αE-catenin to F-actin was also reported in solution [43] and in a biophysical study wherein a single β-catenin/αE-catenin heterodimer formed a short-lived slip bond, but a higher heterodimer surface density enabled the complex to form a directional catch bond with F-actin [35]. Studies from our laboratories likewise indicate that interactions between neighboring cadherin-catenin complexes facilitates F-actin binding under load [18, 30]. These observations suggest that entry into a long-lived binding state may be facilitated by interactions between neighboring complexes. However, to our knowledge how multiple cadherin-catenin complexes interact when under load had not been examined in detail.

To address this question, we first used Monte Carlo simulations based on kinetic parameters derived from the OT experiments to examine how load is distributed when more than one complex is bound to F-actin. For simplicity, we considered two limiting, hypothetical cases: 1) load is shared equally among actin-bound complexes or 2) all of the load is borne by a single complex, where the number of complexes *N* in a discrete cluster ranged from 1 and 5. Contrary to what is experimentally observed, in the equal load sharing model the average binding time per complex decreases (Fig. S5a), because dividing the load among complexes tends to shift all of them into the weak-binding regime that predominates below 5 pN. In contrast, a model in which one complex bears all of the load predicts binding lifetimes that are independent of the number of interacting complexes, which qualitatively matches experimental observations for the ternary complex and for the quaternary complex when load was oriented towards the filament (+) end. In this model, all other bound complexes act as “bystanders” and are subject to no load, undergoing cycles of detachment and rebinding. In reality, “bystander” complexes must necessarily be subject to non-zero load due to their attachment to the filament, but may experience much smaller loads relative to the principal load-bearing complex, such that their dynamic, weak binding interactions with F-actin are unobservable in our assays. Nonetheless, the simplified model in which load is placed on one complex at a time captures the main features of our data.

The fold increase in binding lifetime with step number is roughly constant (Fig. 3A), which is consistent with a model in which load-bearing complex is progressively stabilized by an increasing number of neighboring complexes. As a means of capturing this observation, we developed a model in which neighbor-neighbor interactions lead to increased binding lifetimes (Fig. 4). This model captures the data well (Fig. S6). Remarkably, stabilization of only -1.5 k_B_T per additional bound complex is sufficient to account for the observed increase in binding lifetimes (Fig. S6), indicating that subtle effects can potentially lead to large increases in effective binding lifetimes (see Discussion).

**Fig. 4.**
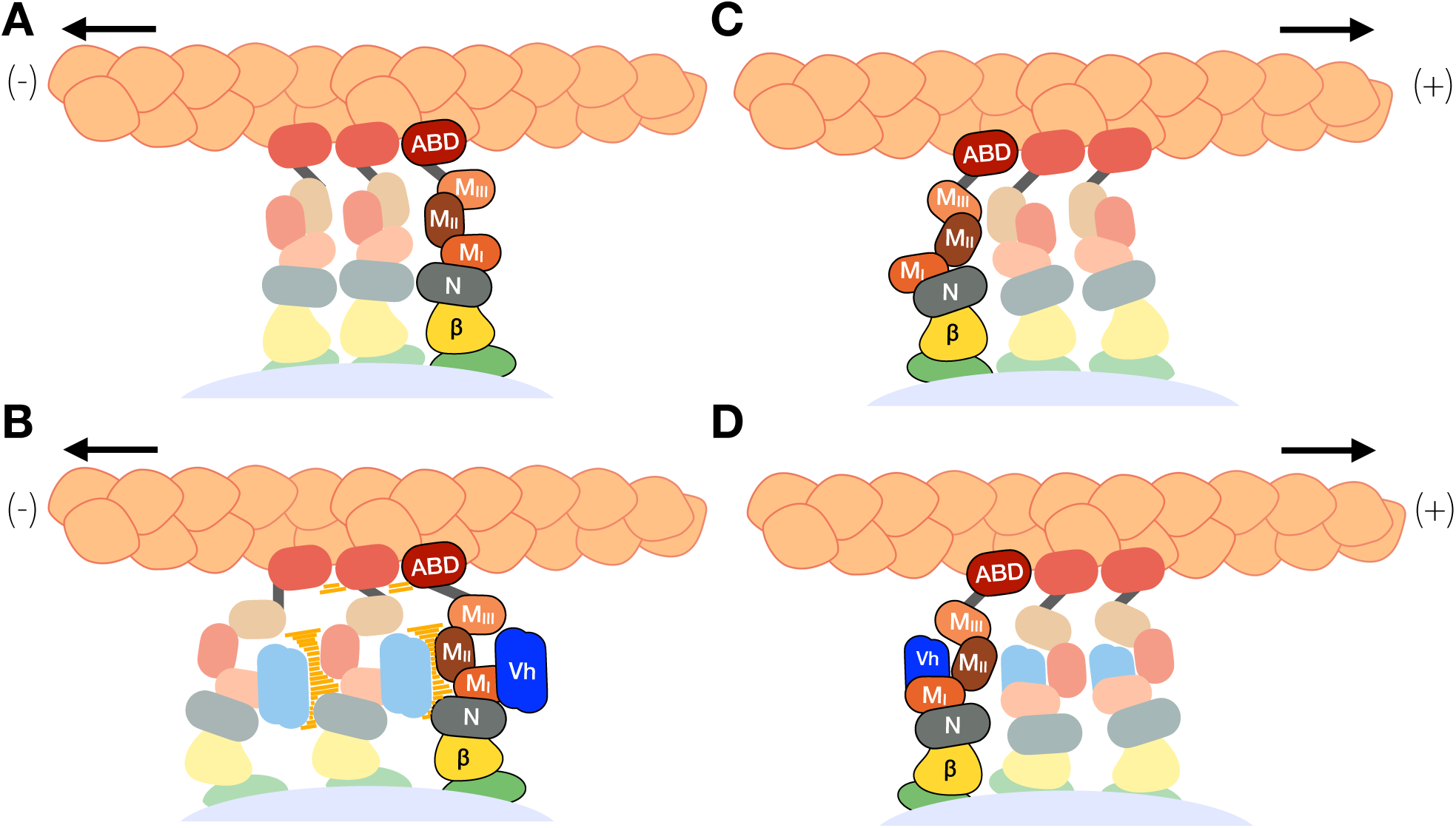
Possible mechanism of increased bound-state lifetimes of cadherin/catenin complexes with (-)-end directed force when vinculin is bound. E-cadherin cytoplasmic domain is shown in green, β-catenin in grey, αE-catenin by its individual domains colored as in Fig. 1, and Vh in purple. The lighter complexes represent bound complexes experiencing no or small load, whereas the darker complexes are experiencing significant load. (**A**, **B**) With force directed in the (-) direction, the M_III_ domain may adopt a position that allows it to contact a vinculin molecule bound to an adjacent complex (orange bars) or cause conformational changes in the ABD. (**C**, **D**) (+)-end directed force produces a different position of M_III_ that cannot contact Vh bound to an adjacent complex or cause conformational changes that would alter binding lifetimes of the neighboring complex.

To further explore the potential ability of a cluster of complexes to cooperatively anchor an actin filament, we calculated the *total time* elapsed until the last of the *N* complexes detached from the filament in Monte Carlo simulations as a measure of force-dependent anchoring. For the ternary complex, total binding times at a given force scaled roughly linearly with *N* (Fig. 5A). This is expected given that our data are best explained by a single complex bearing the large majority of load at any given time (Fig. 5E). In contrast, for the quaternary complex, neighbor-neighbor interactions lead to a large, nonlinear increase in binding lifetimes when load was oriented in the (-) direction (Fig. 5B), consistent with experimental results (Fig. 3). This nonlinearity in turn leads to an asymmetry in binding lifetimes that increases rapidly with *N* for the quaternary, but not ternary, complex (Fig. 5C, D).

**Fig. 5.**
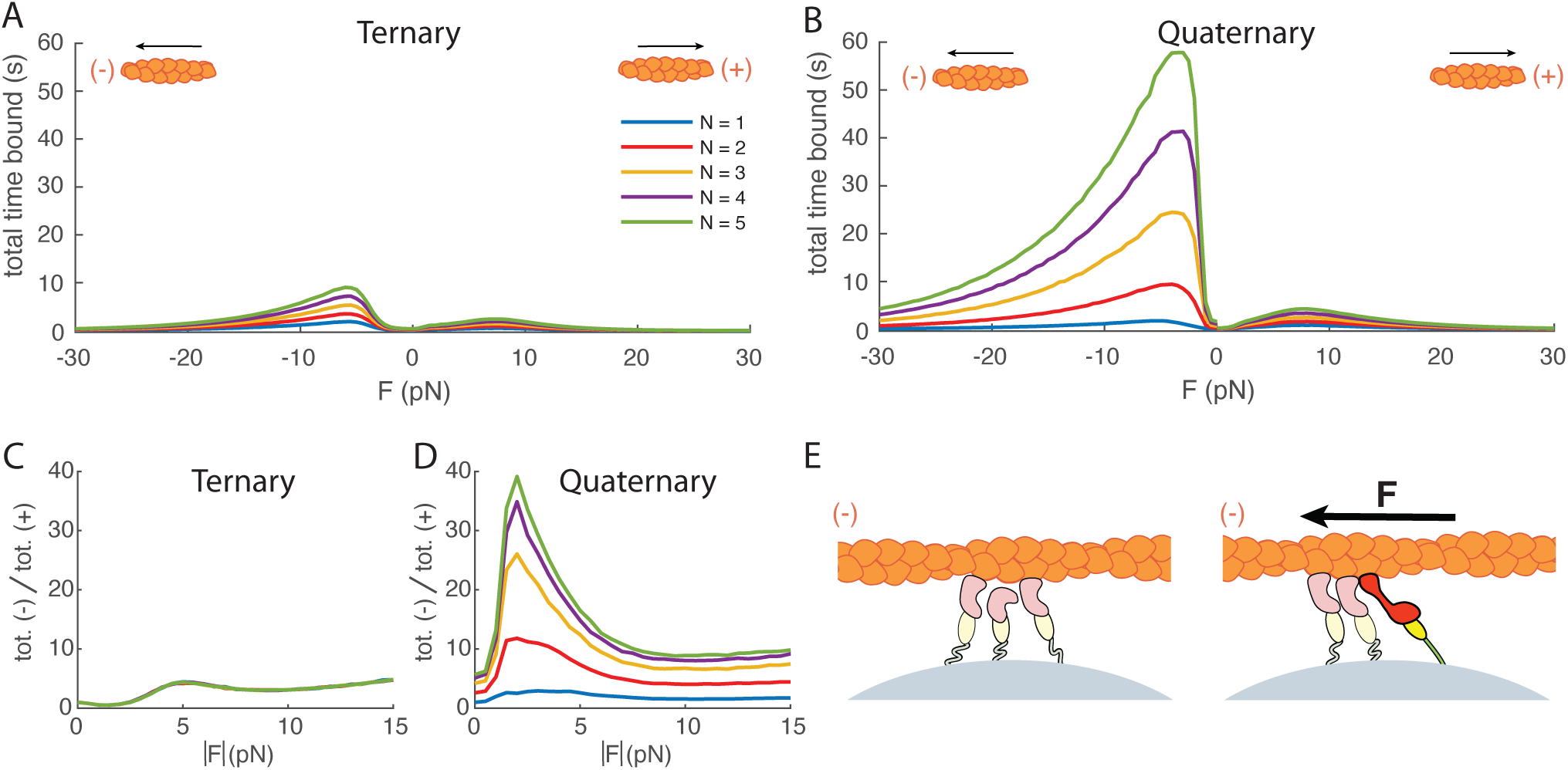
Effects of load sharing and cooperativity on force-dependent actin anchoring. **(A)** Monte Carlo simulation of the total duration until complete detachment for *N* = 1 to 5 ternary complexes, with force ***F*** oriented toward the F-actin (-) end. Detachment is modeled as irreversible. **(B)** Simulation as in (A) but for the quaternary complex. **(C)** Simulated ratio of the time until complete detachment for the ternary complex loaded in the (-) vs. (+) directions. **(D)** Simulation as in (C) but for the quaternary complex. Cooperative interactions between complexes lead to a large, force-dependent increase in directionality. Note that, for the quaternary complex, lifetime ratios are not equal to 1 at zero force due to neighbor-neighbor stabilization when loaded in the (-) but not (+) direction. This represents an approximation of the more physically realistic case in which neighbor-neighbor stabilization may depend on both load direction and magnitude.

Although neighbor-neighbor stabilization is physically plausible, it may not be the only contributor to the directional increase in bound-state lifetimes observed for the quaternary complex. For example, we examined an alternative scenario in which successive change in the angle of the applied force with the number of quaternary complexes interacting with F-actin might alter the degree to which applied load influences the balance between the weak and strong states, and hence overall binding lifetimes (Supplemental Information, Fig. S6, S7). The inherent chirality of both F-actin and the quaternary complex makes it reasonable to suppose that this effect could occur in a direction-sensitive manner. Given their non-exclusivity, alterations in F-actin binding stability and these geometric effects may both contribute to the observed increase in binding lifetime.

## Discussion

We have shown that independent of its own actin-binding activity, vinculin profoundly alters force-dependent binding of multiple cadherin-catenin complexes to F-actin: association of vinculin with the ternary complex of E-cadherin, β-catenin and αE-catenin increases the bound lifetime of individual complexes on F-actin as a function of the number of complexes bound, when force is directed towards the pointed (-) end of the filament. In contrast, the ternary complex does not show this behavior. Thus, force not only promotes both strong binding of the ternary complex to F-actin and to vinculin, but also enables the now-bound vinculin to alter the polarized binding of multiple complexes to F-actin by increasing the mean lifetimes of the individual complexes when force is oriented in the (-) direction. In this way, vinculin may enhance the polarity and stability of the cadherin/catenin/F-actin assembly with load, and thereby reinforce cell-cell contacts.

The molecular mechanism(s) by which vinculin enhances cooperative and directional F-actin anchoring are unclear. It is possible that this arises from favorable interactions between neighboring complexes, reorientation of load in a way that enhances the lifetime of the catch bond, or both (Fig. 4, Fig. S7). Vinculin binding requires unfolding of the αE-catenin M_I_ domain and the loss of interactions that stabilize the relative positions of M_I_, M_II_, and M_III_ [26, 27, 40, 41, 44]. This repositioning of domains may produce new contacts between neighboring complexes that selectively stabilize actin-bound states depending on the orientation of the applied load (Fig. 4B, D). Such neighbor-neighbor stabilization would not occur for ternary complexes, since all but the load-bearing complex would adopt the compact, noninteracting M domain conformation (Fig. 4A, C). Given evidence for allosteric communication between the αE-catenin M-domain and ABD [41], it is possible that the directional repositioning of αE-catenin M subdomains when Vh is bound could allosterically alter ABD conformation and actin binding stability. It is likewise possible that the neighbor-dependent repositioning of αE-catenin domains alters the projection of force along the reaction coordinates that correspond to transitions between states of the ABD catch bond [29], resulting in an increase in binding lifetimes as additional complexes are bound. Experimental tests of these models are, however, beyond the scope of this study.

### Implications of unequal load sharing

When multiple cadherin/catenin complexes bind to an actin filament, it would be reasonable to expect that load would be shared equally among interacting complexes. Instead, we found that bound-state lifetimes for the ternary complex, as well as the quaternary complex when force is oriented in the (+) direction, are most easily explained by a model in which only one complex bears essentially all the load, with the rest acting as bystanders. For the quaternary complex, the increase in mean lifetimes as a function of number of complexes when force is oriented in the (-) direction is also consistent with unequal load sharing, but with an additional source of stabilization that scales with the number of bound complexes. A possible explanation for unequal load sharing is that the force-extension behavior of the cadherin-catenin complex is nonlinear, *i.e.,* more analogous to a rope than a spring. In this view, whichever complex reaches its maximal extension first would bear the majority of the mechanical load. In integrin-based adhesions, a minority of integrins bear the majority of the load [45], suggesting that unequal load-sharing may occur *in vivo*.

Depending on the total load and the force sensitivity of the catch bonds, unequal load sharing could provide a counterintuitive stabilization of the linkage between adhesion complexes and F-actin: one complex bearing most of the load yields nearly constant individual binding lifetimes regardless of the number of complexes bound to F-actin, meaning that how long a filament stays attached to the adhesion complex scales linearly with the number of complexes. If the total force per F-actin filament is similar to the catch bond maximum (∼6 pN for the cadherin-catenin complex), this can produce *longer* total binding lifetimes than equal load sharing, since in the latter case individual binding lifetimes decrease when load is spread among too many complexes (Fig. S5). Consistent with this possibility, both the maximal force generated by nonmuscle myosin II (3.5 pN) [46], and the inferred forces transmitted by individual cytoskeletal linkers in living cells (∼4-8 pN) [45] are comparable to the force at which maximal binding lifetimes occur for the cadherin-catenin [18] and vinculin [29] catch bonds.

### Possible consequences of asymmetric binding to F-actin

Contractile F-actin cables spanning multiple cells power embryonic morphogenesis and wound-healing in epithelia, and muscle contraction in the heart. The actin cables in some epithelial tissues show clear sarcomeric organization, implying that the barbed (+) ends of the filaments terminate at tricellular junctions [13, 15, 47–49] (Fig. 6C). This arrangement is consistent with cell biological, genetic, and electron microscopy data indicating that actin filaments are anchored end-on at epithelial tricellular junctions (e.g. Refs. [50, 51]). Myosin II motor activity is required for the organization of these bundles as well as recruitment of cell-cell junction components [15, 52–56]. An identical molecular-scale organization links myofibrils across the junctions between cardiomyocytes in the heart [57]. However, how these cables can self-assemble to span multiple cells has been unclear.

**Fig. 6.**
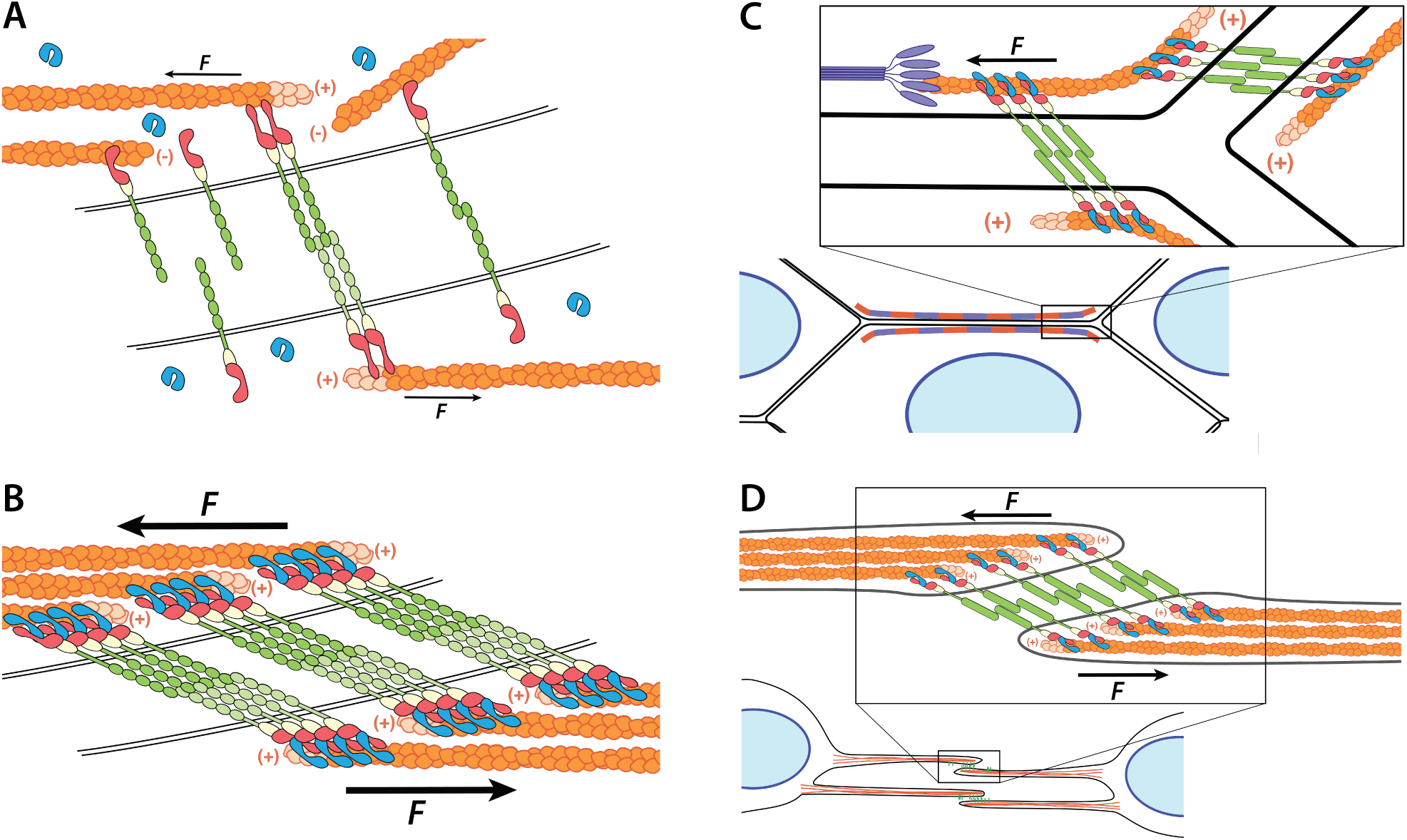
Possible model for assembly of load-bearing connections at cell-cell junctions. **(A)** At low forces, connections between F-actin (*orange*, new (+) ends *light orange*) and the α- catenin/β-catenin/E-cadherin complex (*red*, *yellow*, and *green*) are transient. Vinculin (*blue*) is predominantly in its autoinhibited and cytosolic. (**B**) Force above a threshold opens the vinculin binding site on α-catenin, recruiting vinculin. Cooperative interactions between neighboring quaternary complexes stabilize F-actin loaded toward the (-) end, and simultaneously favor cadherin clustering. **(C)** At tricellular junctions, load-stabilized cadherin-catenin clusters, as in (B), link contractile actin and myosin (*purple*) bundles spanning epithelial tissues. **(D)** Load- driven self-assembly stabilizes and organizes F-actin in contacting protrusions during the assembly of endothelial cell-cell junctions.

We propose that a positive feedback loop stabilizes the connection of the cable at cell-cell junctions (Fig. 6A, B): *i*) Load oriented toward the F-actin (-) end, as generated by nonmuscle myosin II, engages the cadherin-catenin complex catch bond, producing a modest bias in the orientation of the actin filaments. *ii*) Tension on the cadherin-catenin complex leads to the recruitment of vinculin, yielding additional polarization due to the enhancement of binding lifetimes and directionality for multiple, vinculin-bound cadherin-catenin complexes. *iii*) The force-dependent, directional bonds between vinculin and F-actin [29] imparts additional polarization to local filaments. At each step, polarization of F-actin is anticipated to increase the efficiency of force generation by nonmuscle myosin II, leading to a positive feedback loop between myosin contractility, catch bond formation, and F-actin polarization. This feedback loop would result in the ordered sarcomeric assemblies observed in epithelia [13, 15, 58] and cardiomyocytes [57] (Fig. 6C). However, the same feedback loop would be expected to stabilize actomyosin bundles of mixed polarity, though with effectiveness that is correspondingly reduced, given that myosin II can only exert force on filaments oriented with their (+) ends pointing away from the myosin bundle (Fig. 6C). Importantly, myosin II organization appears to precede assembly of mature cell-cell junctions, consistent with the need for tension to promote the polarized binding of αE-catenin and vinculin to F-actin [15, 59].

Directional catch bonds may also play a wider role in driving cell and tissue organization. For example, the organization of F-actin predicted by our model (highly oriented, with barbed ends out) is observed at VE-cadherin based adhesions between the protrusions of neighboring endothelial cells [59, 60] (Fig. 6D). A recent study likewise demonstrates that talin, the principal F-actin binding protein in integrin-based adhesions, also forms a highly directional catch bond with F-actin [61], suggesting a parallel mechanism for the formation of stress fibers to the one explored here. We speculate that directionally polarized binding interactions of the sort described in this study may constitute an important, and presently underexplored, organizing mechanism for the cytoskeleton.

## Methods

### Protein expression and purification

Mouse E-cadherin cytoplasmic domain, E-cadherin cytoplasmic domain aa 785-88, β- catenin, β-catenin 78-671, αE-catenin, αE-catenin R551A, full length chicken vinculin, and chicken vinculin D1 domain (aa 1-259) were purified as described [26, 41]. Vinculin head (Vh; residues 1-851) was expressed with a His_6_ tag in a pET15b vector (kind gift from Dr. Susan Craig) and was purified as previously described [62]. Zebrafish αE-catenin used in the OT experiments was purified as previously described [63].

The expression vector for GFP-E-cadherin used in the OT assay was constructed by inserting DNA encoding the cytoplasmic domain of *Mus musculus* E-cadherin into the pPROEX HtB vector along DNA encoding eGFP to generate an in-frame fusion consisting of an N-terminal His_6_-tag, eGFP, and E-cadherin. GFP-E-cadherin was expressed in BL21(DE3) Codon Plus *E. Coli* cells in LB media at 37 °C. Cells were grown to an OD of 1.0 and induced with 0.5 mM IPTG. After induction, the cells were grown for 16 h at 18 °C, pelleted and resuspended in 20 mM Tris pH 8.0, 150 mM NaCl, 1 mM β-mercaptoethanol and flash frozen. Thawed cell pellets were lysed with an Emulsiflex cell disrupter in the presence of EDTA-free protease inhibitor cocktail set V (EMD Millipore) and Dnase (Sigma Aldrich). The lysate was centrifuged at 27,000 x *g* for 30 minutes. Clarified lysate from 2 L of cells was incubated with 10 mL of TALON Superflow resin (GE Healthcare Life Sciences) for 30 minutes on a rotator at 4 °C. Protein was washed with 5 bed volumes of 20 mM Tris pH 8.0, followed by 5 bed volumes of bed volumes of PBS pH 8.0, 0.5 M NaCl, 0.005% Tween 20, followed by 3 volumes of 20 mM Tris pH 8.0, 150 mM NaCl, 10 mM imidazole, 1 mM β-mercaptoethanol. Protein was eluted from the TALON resin in 20 mM Tris pH 8.0, 150 mM imidazole, 100 mM NaCl, 1 mM β- mercaptoethanol. The eluate was passed through a 0.22 µm SFCA syringe filter and diluted to a final volume of 50 mL in 20 mM Tris pH 8.0, 1 mM DTT, 0.5, mM EDTA. The filtered eluate was applied to a MonoQ anion exchange column in 20 mM Tris, pH 8.0, 1 mM DTT and run with a 0-1 M NaCl gradient and protein eluted at approximately 300 mM NaCl.

Proteins were stored at -80°C and never underwent more than one freeze/thaw cycle.

### Actin cosedimentation assay

Rabbit muscle G-actin was polymerized by addition of 10x polymerization buffer (500 mM KCl, 20 mM MgCl_2_ and 10 mM ATP in 100 mM Tris, pH 7.5) and subsequent incubation at room temperature for 1h. Ternary and quaternary complexes were formed by mixing αE-catenin with excess E-cadherin cytoplasmic domain, β-catenin and vinculin Vh or D1. After incubation for 15 min on ice complexes were purified on a S200 gel filtration column (GE healthcare) equilibrated with 10 mM HEPES pH 7.5, 50 mM KCl, 1mM DTT and 2 mM MgCl_2_. For binding assays a concentration series ranging between 15 μM and 0.25 μM was generated by serial dilution into assay buffer (10 mM HEPES pH 7.5, 50 mM KCl, 1 mM DTT and 2 mM MgCl_2_). An equal volume of 3 μM F-actin was added (final F-actin concentration 1.5 μM) and incubated for 30 min at room temperature. For the ternary complex with αE-catenin R551A, the assay was performed in 100 mM KCl due to solubility limitations of the complex. To determine nonspecific background precipitation an identical concentration series was prepared with F-buffer (5 mM Tris-HCl pH 8.0, 0.2 mM CaCl_2_, 50 mM KCl, 2 mM MgCl_2_, 1 mM ATP, and 0.5 mM DTT) instead of F-actin. Actin-bound αE-catenin was separated by centrifugation at 55,000 rpm for 20 min at 20 °C in a TLA 100 rotor (Beckman Coulter, Inc.). The F-actin pellet was re-suspended in sample buffer and run on SDS-PAGE. Coomassie-stained bands were imaged and analyzed with a LI-COR Odyssey imaging system and the LI-COR analysis software (LI-COR Biotechnology). Background was subtracted for each concentration point and the amount of pelleted αE-catenin was normalized to the amount of pelleted actin. Concentrations were extrapolated from a standard curve. Data were analyzed with the Prism analysis software (GraphPad Software, Inc.). Note that at high concentrations there is significant background sedimentation of either αE-catenin or the ternary complex (Fig. S4), making precise quantification of these assays difficult, so we cannot not rule out a small effect of the R551A mutation or vinculin on actin binding in solution.

### Optical Trap Assay

In this assay a biotinylated actin filament links two optically-trapped, streptavidin-coated beads to create a “dumbbell” (Fig. 2A). The actin filament is then positioned over a surface-immobilized “platform” bead bearing cadherin/β-catenin/αE-catenin complexes. The microscope stage is moved in a trapezoid-wave pattern, such that the binding of complexes on the platform bead results in the displacement of one of the two optically trapped beads (Fig. 2A). The stage motion stops if a binding event is detected at the end of a 5 ms loading phase.

When a displacement is detected, the stage motion halts and the displaced bead relaxes back to its equilibrium position as the bound complexes release from the filament. The last release step corresponds to the dissociation time of a single complex. Because the optical trap acts as a spring with a known stiffness, the displacement provides the force exerted on the bead. Once both optically trapped beads return to their baseline position, the stage motion resumes, allowing us to record multiple such binding events per platform bead.

The optical trap assay was carried out as described [18], with 50 μM GFP-E-cadherin cytoplasmic domain, 100 nM β-catenin, and 75 nM αE-catenin, except the final buffer injection also included 1 μM Trolox (Sigma Aldrich). Zebrafish αE-catenin was used in these experiments to ensure that only monomeric αE-catenin was added [18]. For vinculin experiments, 15 μM of Vh was added in the final injection. This concentration was chosen based on the K_D_ of 1.9 μM of the vinculin D1 domain for the ternary complex in solution [26], which implies that approximately 90% of the complexes would be bound to vinculin D1 in the absence of force. We were unable to obtain a direct ITC measurement of the affinity of Vh for the ternary complex, likely because the enthalpy change is very small, as found for D1 [26], but since both Vh and D1 bind to the minimal vinculin-binding fragment of αE-catenin with similar affinities [26], we assume that their affinities for the wild-type complex are comparable. Importantly, force promotes binding of vinculin to αE-catenin [27], so the effective K_D_ in the OT experiments is likely to be higher. Control experiments in which Vh or buffer alone was added to the flow cell containing beads bearing cadherin cytoplasmic domain and β-catenin, but no αE- catenin, showed no significant binding to F-actin.

Every binding interaction which survived the 5 ms load phase was included. We used the previously described directionality assay [29] to determine the polarity of a subset of the actin filaments, and used this subset to infer the directionality of all of the filaments that had sufficient data to be statistically significant (see SI text). Modeling of the data was constrained such that the mean lifetime at zero force was less than or equal to the mean lifetime measured using a low-force OT binding assay, as previously described [29].

### Selection of two-state catch bond model and statistical analysis

To model the OT data, one-state slip and catch bond models, as well as a two-state slip bond model, described previously [18, 29], were considered. One-state models were ruled out because they could not capture the biexponential distribution of lifetimes at a given force. The two-state slip bond model was ruled out because it cannot describe the biphasic behavior of the force-lifetime curve. Details of the two-state directional catch bond model, statistical analysis and parameters are provided in Supplementary information and Table S1.

### Simulations

Details of the Monte Carlo simulations are provided in the SI.

## Acknowledgements

We thank Dr. Craig Buckley for his contributions to the early stages of this project. Research reported in this publication was supported by a Howard Hughes Medical Institute Faculty Scholar Award (A.R.D), as well as National Institutes of Health grant R01GM114462 to W.I.W. and A.R.D, R35GM130332 to A.R.D. and R35GM131747 to W.I.W. N.A.B. and D.L.H. were supported by training grant T32 GM007276 from the NIH. A.W. and D.L.H. were supported by Graduate Fellowships from the National Science Foundation. A.W. was also supported by a Stanford Graduate Fellowship and training grant T32GM120007. Use of the Stanford Synchrotron Radiation Lightsource, SLAC National Accelerator Laboratory, is supported by the U.S. Department of Energy, Office of Science, Office of Basic Energy Sciences under Contract No. DE-AC02-76SF00515. The SSRL Structural Molecular Biology Program is supported by the Department of Energy Office of Biological and Environmental Research and by the National Institutes of Health, NIGMS Grant P30GM133894. The contents of this publication are solely the responsibility of the authors and do not necessarily represent the official views of NIGMS or NIH.

## Data availability

The OT data, analysis scripts and modeling computer code used in this work are available upon reasonable request from the corresponding authors.

## Supplemental Information

### Determination of actin filament polarity

Because the myosin VI directionality assay (1) has a relatively low throughput, it was only used to determine the directionality of a subset of actin filaments used in our study. The data from filaments with known polarity were used to make a reference set. We then used a statistical test to determine whether filaments without experimentally determined polarity had sufficient binding events to confidently infer their polarity. Data from the reference set was separated into four categories: 1) single-step events with load directed towards the pointed (-) end of F-actin, 2) multi-step events with force directed towards the pointed (-) end of F-actin, 3) single-step events with force directed towards the barbed (+) end of F-actin, and 4) multi-step events with force directed towards the barbed (+) end of F-actin. For directionally dependent observations, we will henceforth use superscript notation to denote the polarity of the actin filament. For instance, ‘quaternary^(-)^ complex’ will refer to quaternary complex loaded in the (-) direction. For each filament whose polarity was not determined experimentally, we assigned a provisional polarity based on the sum of lifetimes measured for a given binding direction (*i.e.*, force towards trap 1 or trap 2). We then counted the number of single- and multi-step events in each direction, and generated 10,000 synthetic datasets with the same number of events in each of the four categories by drawing random events, with replacement, from the known polarity reference dataset. Next, we computed the inferred directionality for these synthetic datasets: in cases where the total number of events was sufficiently small, the inferred orientation for a given, synthetic dataset was sometimes ‘flipped’ relative to the known polarity due to the stochastic sampling of the reference dataset. We thus included datasets corresponding to filaments with a sufficient number of binding events such that the corresponding, simulated datasets flipped directionality in less that 1% of trials.

### Fit of last-step data to a directional two-state catch bond model

As in previous studies, we reasoned that the last observable plateau prior to complete detachment from F-actin reflected the binding lifetime of a single cadherin-catenin complex. These binding lifetimes are empirically well-described by a biexponential function across the large majority of the force range probed in our experiment. This observation is consistent with the presence of (at least) two distinct F-actin bound states (2). The data were thus fit to a two-state directional catch bond, as described previously (1).

To fit the force-lifetime measurements for both the ternary and quaternary complexes, we approximated the increase in load that occurs when the previous complex detaches as essentially instantaneous. This assumption is reasonable provided that the timescale for relaxation of beads in the optical trap (∼1 ms) is smaller than the timescale for of the interconversion between F-actin bound states and/or detachment. Accordingly, we used a constant flux condition (2, 3) to ensure that the model obeyed the principle of detailed balance and to calculate the ratio of rates for binding into the weak or strong states (k_01_ and k_02_ respectively) (Eq. 1).

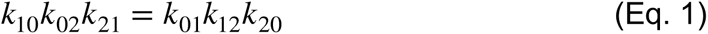

The two-state directional catch bond model contains 12 adjustable parameters, *i.e*. four rate constants and eight distance parameters (1). We fixed *x_10_^(-)^* and *x_10_^(+)^* (the distance parameter in the negative and positive directions) at 0 nm, because they were well-constrained at < 0.1 nm in a coarse-grained parameter search (*not shown*). In addition, *x_21_^(-)^* and *x_21_^(+)^* were in general not well constrained, reflecting the fact that the output of the model depended more strongly on the equilibrium between the strong and weak states, *i.e., k_12_*exp(*x_12_F*/*k_B_T*) / *k_21_*exp(-*x_21_F*/*k_B_T*), than on any one of the four underlying parameters. In particular, we found that global minimization yielded poorly constrained values for *x_21_^(+)^* for the ternary cadherin-catenin complex, and *x_21_^(-)^* for the quaternary cadherin-catenin-Vh complex. To avoid these parameters converging to unrealistically large values, we fixed these two parameters at -15 nm, a value that represents a plausible upper bound for the extension that H0 and H1 undergo when undocked from the rest of the ABD under load (4). These approximations reduced the model from 12 to 9 adjustable parameters (Table S1).

We used a maximum likelihood framework to fit the 9-parameter model to the set of individual force-lifetime measurements for both the ternary and quaternary complexes. As in our previous study, we used the binding lifetimes measured at close-to-zero force to constrain the model (1). MLE fitting was performed using two strategies: In one approach, we used a constrained nonlinear convex optimization routine (Matlab: `fmincon`) to minimize the negative log likelihood score, seeding with 10^5^ – 10^7^ starting points generated by the Matlab `Multistart` routine. In an alternative approach, we minimized the negative log likelihood score using the Matlab implementation of the genetic algorithm (Matlab: `ga`). 10-100 epochs of the genetic algorithm were required to reliably identify a global minimum. Both approaches were comparably successful at finding global minima. However, in our hands the genetic algorithm was considerably faster. It was therefore used for bootstrapping model confidence intervals (see below).

### Estimation of parameter confidence intervals (CIs)

We used profile likelihood as a means to estimate the confidence intervals for individual parameters (5). Briefly, the upper and lower bounds for a given parameter are determined as follows: A given parameter, generically *θ*, is systematically varied about its optimal value, while all other parameters are optimized for each value of *θ.* The resulting log-likelihood ratios asymptotically approach a *χ*^2^ distribution with one degree of freedom. The upper and lower bounds are thus determined by Eq. 2:

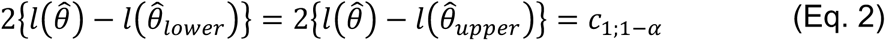

Where *l* is the log-likelihood function, *θ̂* denotes the MLE for *θ*, <u>and</u> *c*_1;1-*α*_ is the (1-α)th quantile of the *χ*^2^ distribution with 1 degree of freedom. 5^th^ and 95^th^ percentile bounds for each parameter were determined by a grid search followed by linear interpolation (Table S1).

### Model confidence intervals (CIs) and model comparisons

In general, it is not straightforward to go from parameter CIs (above) to CIs for the resulting model. Thus, we determined CIs for the model for both the ternary and quaternary complexes using a Bootstrap approach, as described previously (1, 2, 6). Briefly, synthetic datasets were constructed by drawing, with replacement, *n* force-lifetime observations (the same number as in the original dataset). These datasets were then fit using the genetic algorithm, as above. A small fraction of these fits (0.4% for the ternary complex, 0.7% for the quaternary complex) did not converge to models that obeyed the low-force lifetime constraint. These fits were excluded from the subsequent analysis. As a result, CIs were constructed from 1985 synthetic datasets for the ternary complex, and 996 synthetic datasets for the quaternary complex.

To determine whether the models for the ternary and quaternary complexes differ from each other in a statistically meaningful way, we used a version of bootstrap hypothesis testing (6). Specifically, we drew at random 10^6^ pairs of bootstrapped parameter sets for the ternary and quaternary complexes. We then calculated the fraction of the resulting families of models, **M_T_**(*F*) and **M_Q_**(*F*), respectively, for which the average lifetime of the quaternary complex was longer than that of the ternary complex at a given force *F*. We ascribe statistical significance when the fraction **M_T_**(*F*) > **M_Q_**(*F*) is > 0.975 or < 0.025 for a given *F* (Fig. S3).

### Determination of binding lifetimes at low force

We sought to characterize the binding lifetimes of the ternary cadherin-catenin complex and the quaternary cadherin-catenin-Vh complex at close to zero load. We repeated the standard optical trap assay but with a peak-to-peak oscillation of 40 nm and without halting the stage upon binding. As a result, the magnitude of the maximal force exerted on either optically trapped bead was ∼4 pN, with most events at lower forces. Binding events measured in this manner likely represent an upper bound on the true lifetime at zero force due to a likely modest increase in binding lifetimes at low, non-zero forces. We cannot exclude the possibility that some of the binding events we observe in this assay reflect the simultaneous binding (and release) of more than one complex, which may also inflate the apparent binding lifetime. Mean binding lifetimes were 10 ms (control; N = 35), 62 ms (ternary complex; N = 90) and 44 ms (quaternary complex; N = 210) (Table S3).

Even with optimized conditions, occasional transient interactions between the actin filament and platform bead were observed in a control measurement in which only GFP-E-cadherin cytoplasmic domain, β-catenin, and Vh were added (*i.e*. no αE-catenin). We used the following strategy to account for the contribution of these non-specific background binding events in the ternary and quaternary datasets: First, we noted that the cumulative distribution for lifetimes measured for the control sample was empirically well described by a biexponential function with rate constants of 180 and 40 s^-1^, with the fast phase accounting for 78% of the total observations. We therefore excluded events shorter than 20 ms from the analyses of the ternary and quaternary datasets, as this cutoff should reject ∼97% of the short-lived background events, and 88% of background events overall. Both the ternary and quaternary datasets were well described by biexponential functions, with rate constants of 30 and 2.5 s^-1^, and 37 and 1.8 s^-1^, respectively. For the ternary complex, extrapolating back to time *t* = 0 predicted 66 binding events corresponding to the 30 s^-1^ rate constant, and 8 events corresponding to the 2.5 s^-1^ rate constant. A similar calculation for the quaternary complex yielded a predicted 151 and 9 events corresponding to the faster and slower rate constants, respectively. Comparison with the actual number of measured events indicated the presence of 16 and 50 “excess” events for the ternary and quaternary complexes, respectively, presumably reflecting nonspecific interactions detaching at ∼180 s^-1^ that were excluded by the 20 ms cutoff.

The 40 s^-1^ rate constant for detachment observed for the control sample is, in practical terms, not possible to distinguish from the 30 and 37 s^-1^ rate constants for the ternary and quaternary complexes, respectively. To account for the contribution of this minor population of non-specific interactions to the observed kinetics, we first took the calculated “excess” events, above, to be fast-detaching (180 s^-1^) nonspecific interactions. We then inferred the corresponding number of intermediate rate (40 s^-1^), nonspecific events, which should be ∼22% the number of fast-detaching nonspecific binding events. This predicted 4 and 14 background events occurring at ∼40 s^-1^ for the ternary and quaternary complexes, respectively. For the ternary complex, subtracting these counts yielded 62 “true” events corresponding to the 30 s^-1^ rate constant, while a similar calculation for the quaternary complex yielded 137 “true” events corresponding to the 37 s^-1^ detachment rate constant. Combined with events corresponding to the slow detachment rate (2.5 s^-1^ and 1.8 s^-1^, ternary and quaternary complexes), these values yielded predicted binding lifetimes of 76 and 60 ms for the ternary and quaternary complexes, respectively. These values represent ∼5% corrections relative to the lifetimes predicted from biexponential fits described in the previous paragraph.

### Multistep data

We define “binding event” or “event” as the period of time during which the actin filament is connected to the platform bead through one or more ternary or quaternary complexes. To avoid repetition, we refer to steps with reference to the end of a binding event, *i.e*. ‘first’ denotes first-from-end (*i.e*., last), ‘second’ denotes second-from-last step, and so on.

### Statistical comparison of binding lifetimes as a function of step number

We first sought to determine whether the lifetimes of a given step number differed in a statistically significant way for the ternary versus quaternary complexes. Accordingly, we used a two-dimensional Kolmogorov–Smirnov test to test the hypothesis that the distributions of binding lifetimes for a given step number were drawn from the same underlying distribution, with the threshold for statistical significance subject to a Bonferroni correction for 5 comparisons (Table S4). By this standard, the 1^st^ through 4^th^ steps found to differ significantly for the ternary and quaternary complexes loaded in both the (+) and (-) directions. The 5^th^ steps did not differ in a statistically significant way, likely due to the limited number of observations.

### Force dependence of average binding lifetimes for multiple complexes

We used boxcar averaging to calculate the average binding lifetime for events within a sliding 2.5 pN window spaced every 0.25 pN. We then calculated the ratio of average binding lifetimes for 2^nd^ vs. 1^st^, 3^rd^ vs. 1^st^, and 4^th^ vs. 1^st^ steps (Fig. S8). (The limited number of observations for the 5^th^ step did not allow this calculation.) For the quaternary^(-)^ complex, this analysis did not yield an obvious trend in the enhancement of binding lifetimes as a function of load. This observation is consistent with the idea that the presence of multiple bound complexes increases the stability of the load-bearing complex, leading to a decrease in *k_10_^(-)^, k_20_^(-)^* or both. However, the averaged data are sufficiently variable that we cannot exclude an additional alteration in *x_20_^(-)^* or other distance parameters.

### Binding lifetime as a function of the number of interacting complexes

We calculated the average bound-state lifetime across all forces for the 1^st^ through 5^th^ steps for both the ternary and quaternary complexes with (+) and (-)-end directed loads (Table S2, Fig. 3). In doing so, we noted an increase in binding lifetime with step number for the quaternary^(-)^ complex. We sought a function with the minimum number of parameters that could empirically describe the increase in binding lifetimes observed for the quaternary^(-)^ complex. An exponential function of the form *L*(*n*) = *L*_1_*exp*(*c*·(*n*-1)), where *L*(*n*) is the expected lifetime for a step number *n*, *L_1_* is the lifetime of the first-from-end step (*i.e*., last step), and *c* a free parameter, described the data well. Fig. 3 shows the weighted, nonlinear least squares fits, where the weights are the reciprocal of the variance on the bootstrapped mean for each step number.

We next used bootstrap fitting to test the hypothesis that the trends in binding lifetime vs. step number differed in a statistically meaningful way. We sampled events for steps one through five with replacement and fit each of these synthetic replicates to the exponential function above. The resulting values of *c* for the quaternary^(-)^ complex differed significantly from zero (Table S5). Interestingly, the same held true for the ternary^(-)^ complex, as binding lifetimes *decreased*, rather than increased, with step. The bootstrap fitting as described above was used to calculate the 95% confidence intervals shown in Fig. 3.

### Kinetic Monte Carlo simulations of cadherin-catenin clusters bound to F-actin

We used kinetic Monte Carlo simulations to investigate possible mechanisms by which Vh might modulate the cooperative binding of the cadherin-catenin complex to F-actin. In the models described below, we consider *N* complexes initialized in either the weak or strong state F-actin binding states. In our treatment, molecules were initialized in either the strongly or weakly bound state according to the principle of detailed balance (Eq. 1). To estimate an effective *k_on_*, we noted that the dissociation constant under no load for the ternary and quaternary complexes to F-actin are both ∼10 μM (Fig. 1E), and, as measured here, the dissociation rate at zero load is ∼17 s^-1^, indicating a second-order *k_on_* of 1.7 · 10^6^, M^-1^ s^-1^. In addition, the length of the cadherin-catenin complex is ∼22 nm (Fig. S9). Assuming that the surface-attached cadherin-catenin complex explores a half-sphere, an estimated first-order on-rate is then ∼100 s^-1^, although the actual value may differ by several orders of magnitude.

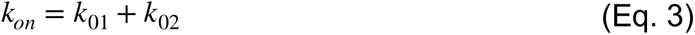

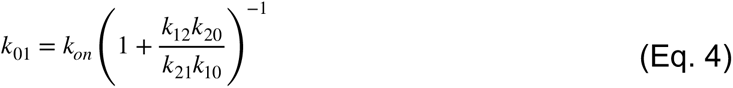

### Bond survival time for complexes sharing equal load

We first consider the case in which the load is shared equally among all bound complexes. This scenario can be treated analytically using a bond survival probability, *B(t)*, derived previously (2, 3) (Eq. 5). From this result, we arrive at a cumulative distribution function (CDF) for the likelihood of *not* observing detachment of the *N^th^* complex in time *t* (Eq. 6). From the CDF, the mean lifetime of the N^th^ step, 〈*τ*〉*_N_*, can then be calculated from numerical integration (Eq. 7, 8).

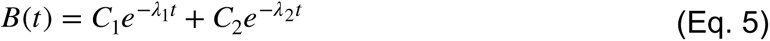

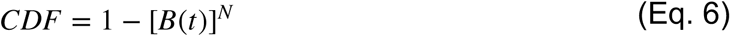

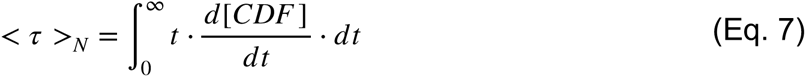

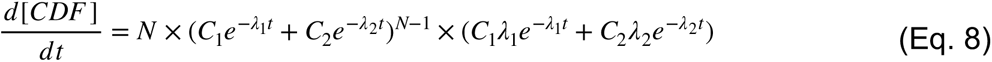

The assumption that bound complexes share equal load results in the prediction that 〈*τ*〉*_N_* should decrease as more complexes are bound to F-actin, which contradicts the experimental observations (Fig. S5).

### Bond survival time for complexes with unequal load sharing

We next consider a limiting model in which one complex bears all of the load. This is a simplification of the real case, in which load is likely to be unevenly distributed amongst bound complexes. However, a model in which load is evenly distributed does not describe the data (see above). Instead, a model in which the large majority of the load is borne by one complex at a time successfully describes the results observed for the ternary complex with load oriented in either direction, as well as the quaternary complex when load is oriented in the (+)-end direction. Uneven load sharing is physically plausible: Given the geometry of the assay, it is likely that load will be distributed unevenly among bound complexes (Fig. 5E). In addition, for forces below 2 pN, both the ternary and quaternary complexes are expected to rapidly equilibrate between bound and unbound states, an effect that is anticipated to concentrate load on one strongly bound complex. Equilibration of the non-load-bearing complexes is also consistent with the description of last steps as starting from an initial condition based on flux balance (see *Fit of last-step data to a directional two-state catch bond model*, above). We note that transient binding/unbinding events of this sort would be difficult to detect in our assay, as they correspond to changes in force that are below our noise threshold.

To avoid introducing additional parameters, we captured unequal load sharing in a model in which all but one interacting complex are assumed to experience negligible force until rupture of the initially loaded complex, when load shifts to another, randomly selected bound complex. For simplicity, the load is kept constant even as individual complexes detach. In developing the model, we noted that the distribution of the number of steps per binding event is roughly power law distributed (Fig. S10), with 7 being the largest number of steps that were experimentally observed in any dataset. Thus, it is likely that, under the conditions of our assay, the maximum number of complexes bound to F-actin is relatively small. In addition, the magnitude of the load decreases between steps, while variable, was typically ∼2 pN, corresponding to ∼20 nm steps at the combined trap stiffnesses used in this experiment. Finally, the cadherin-catenin complex binds to F-actin in a cooperative manner (2, 7). These latter two considerations suggest that the complexes bind in close proximity. Finally, our data are most easily accounted for by a model in which the detachment of the loaded complex is irreversible, as this provides a straightforward means to account for the experimentally observed power-law distribution in the number of steps per binding event (Fig. S10). One physical explanation for this effect is that when a load-bearing complex detaches, the geometry of load application may shift such that the complex can no longer access the filament.

Thus, to model *i*) the experimentally observed distributions of number of steps per binding event and *ii*) the step duration as a function of step number, we initialized simulations with a variable number of complexes up to a maximum number *N* (in the implementation here, *N* = 7). This value reflects the maximum number of steps observed for a single binding event, and also corresponds to the approximate number of accessible binding sites along one face of a F-actin pseudohelical repeat. The number of complexes at the beginning of the simulation is assigned by considering a 1D lattice of potential attachment sites for the quaternary complex on the surface of the platform bead, with an occupancy probability *p_O_* at each site of 0.15. This results in an approximately Poisson-distributed number of complexes that can access the filament, which can account for the approximately power-law distribution in the number of steps per binding interaction. Although it likely represents a simplification, we modeled each complex as interacting with only one site on F-actin, *i.e*., if a given complex dissociates it can only rebind to the same site. At the start of the simulation, one bound complex is assigned to bear all the load. When it detaches, load is transferred to another randomly selected bound complex. The simulation continues until all complexes have detached.

To account for the increase in binding lifetime with step number observed for the quaternary^(-)^ complex, we include an energetically favorable neighbor-neighbor interaction ΔE between immediately adjacent complexes (Eq. 9). Given the linear nature of the actin filament, each complex can be stabilized by interactions with at most *n* = 2 neighboring complexes. For the quaternary^(-)^ complex dataset, the fold increase in binding lifetime with step number is roughly constant regardless of step number (Fig. S8). For this reason, and for parsimony, we assume that negative ΔE stabilizes the weak and strong actin binding states to the same extent. In principle this could influence both binding and detachment rates (*e.g. k_01_* and *k_10_*, *k_02_* and *k_20_*). In the implementation shown here, we model ΔE as altering *k_10_* and *k_20_* only (Eq. 9). This choice embodies the physically reasonable assumption that the transition states are closer in conformation to the bound states than the unbound state.

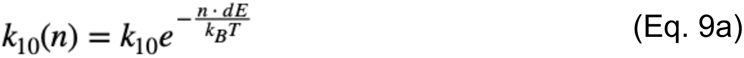

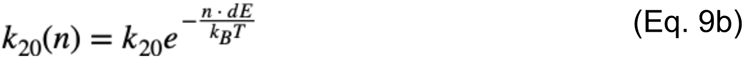

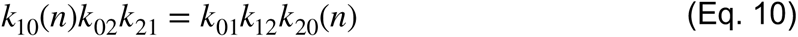

With these assumptions in place, simulations are performed as follows: The number and composition of clusters are defined as above. Individual complexes within the clusters are initially assigned to the weak, strong, or unbound state according to detailed balance at zero load (Eq. 10). Next, one of the bound complexes is randomly selected as the load-bearing complex. The time evolution of the system is performed using the Gillespie algorithm (8) until a timestep occurs in which no complexes are bound to F-actin, meaning that the filament has detached. To develop statistics, the simulation was run 10^6^ times at 8 pN. Two parameters, *p_O_*, and ΔE, were empirically tuned to replicate the distribution of steps per binding event and the increase in binding lifetime with step number in the quaternary^(-)^ complex dataset (Fig. 3).

We were interested to examine how cluster size might affect F-actin polarization as a function of load (Fig. 5). We initialized simulations as above, but with *N* = 1, 2, … 5 complexes and *p_o_* = 1, while systematically varying force *F* from -30 to 30 pN. For each simulation, we calculated the time until all of the complexes were simultaneously detached from the F-actin filament (*i.e*. total bound time). The simulation was repeated 10^4^ times for each pair of *N* and *F*, and the results averaged to yield the plots in Fig. 5A- D.

### Geometric model

The neighbor-neighbor stabilization model above is not unique in its ability to explain the qualitative features of the data. As a non-exclusive alternative, we considered a model in which the binding of additional complexes to F-actin affects how force modulates the projection of the force vector *F* onto *r_ij_* (Eq. 11) (Fig. S6, S7).

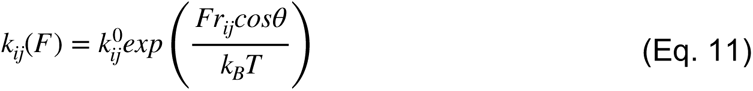

Specifically, we consider the scenario where the binding of additional complexes to F- actin changes %, the alignment between force *F* and vector *r_ij_*, which alters the apparent *x_ij_* (Eq. 12). We make the simplifying assumption that each additional bound complex changes *x_ij_* by a constant value *Δx_ij_*, such that a linear relationship exists between *x_ij_* and the number of bound molecules, *N* (Eq. 13, 14). Such a relationship predicts an exponential dependence on *k_ij_* on *N* for a given force, in empirical agreement with observation for the quaternary^(-)^ dataset (Fig. S6).

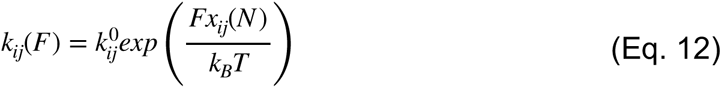

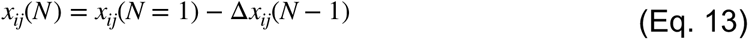

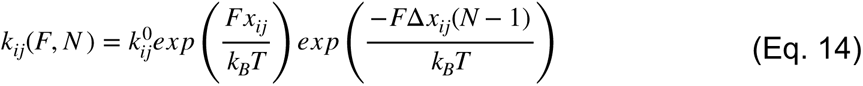

To investigate the impact of *x_ij_* modulation on the two load-sharing models described in earlier sections, we simulated 1,000 parameter combinations in which *Δx_10_, Δx_20_, Δx_12_, Δx_21_* were simultaneously uniformly sampled for each condition, where *Δx_ij_ ∈* [-1,1] nm (Fig. S11). Predicted lifetimes at 8 pN were averaged over 5000 runs for each parameter set sampled.

We were interested to reexamine the question of equal vs. unequal load sharing in the context of this particular model. To do so, we performed the parameter sweep above, but with equal or unequal load sharing. Every set of parameters sampled with evenly distributed load between complexes predicted an inverse relationship between bound lifetimes and *N*, with mean lifetimes consistent to Fig. S5A. In contrast, of the parameters sampled for the one-load bearing complex model, 38.6% resulted in a positive correlation between bond lifetimes and *N*, 32.7% resulted in decreased correlation between lifetimes vs *N*, and the remaining conditions did not predict discernible trends in lifetimes with respect to complexes bound. All parameter sets examined in which *Δx_20_* < 0 resulted in a pronounced increase of binding lifetimes with respect to number of complexes, where *Δx_20_* = -0.27 nm predicts lifetimes consistent with experimental observations (Fig. S5B). Thus, exponentially scaling *k_20_(F)* by number of bound complexes in the unequal load sharing model is sufficient for modeling the dependence of binding lifetimes on *N* (Fig. S6, S7).

We note that although the magnitudes of *x_12_* and *x_21_* are larger than *x_20_*, additional simulations with 1 < |*Δx_21_* |, |*Δx_12_* | < 20 still did not materially affect the relationship between bound lifetimes and *N* for F = 8 pN. The results from our simulations suggest that a minimal model in which the alignment between *F* and *r_20_* decreases as *N* increases suffices to describe the cooperativity experimentally observed when Vh is present.

A weakness of this model is that while *x_20_* and *Δx_20_* can be estimated, *r_20_* and *θ_20_* are not directly measurable from the kinetic data. However, we can place some constraints: *r_20_* ≥ *x_20_* by definition, and the magnitude of *r_20_* is constrained by the plausible size of the underlying structural transition in the protein during unbinding (2). In addition, the change in *θ_20_*, Δ*θ_20_*, with *N* is likely to be small, since it is physically implausible that the F-actin filament will rotate more than, for example, 90° when 8 complexes bind along one face of the pseudohelical repeat (i.e., ∼11° per complex, maximum). Assuming values of r_20_ = 4 nm and Δr_20_ = 5° was sufficient to account for the increase in binding lifetimes with step number for the quaternary^(-)^ complex dataset.

### Small Angle X-ray Scattering (SAXS) analysis of the ternary aE-catenin/b- catenin/E-cadherin complex

For SAXS analysis, a freshly purified monomeric fraction of full-length murine αE- catenin was incubated for 5 min at room temperature with a molar excess of β-catenin 78-671 and a construct comprising the C-terminal 106 amino acids of E-cadherin. The ternary complex was purified on a S200 gel filtration column and the peak fraction was concentrated to 21 mg/ml.

Size exclusion chromatography (SEC) coupled SAXS data were collected at beamline 4-2 at the Stanford Synchroton Radiation Lightsource. Runs comprising 30 and 50 μl of 11 mg/ml or 50 μl of 21 mg/ml were injected onto a 2.4 ml SEC column (Superdex PC 3.2/300). The column was equilibrated with 20 mM HEPES pH 8.0, 150 mM NaCl and 5 mM DTT at a flow rate of 0.075 ml/min. The eluate was directed through a quartz capillary and scattering data were recorded in 1 s intervals on a Pilatus 3XIM detector with a 0.3 x 0.3 mm beamsize and a 1.7 m detector distance. Data processing was performed at the beamline with SasTool. For buffer subtraction, the first 100 images were averaged and used as a buffer profile. The automatically calculated R_g_ is plotted together with the extrapolated I(0) value. Frames from the center of the peak (frame nos. 320-324) were averaged and scattering curves from the 3 data sets were scaled and merged for analysis. The radius of gyration was determined from the linear region of the Guinier plot with an sR_g_ limit < 1.3 using the program Scatter (Robert P. Rambo). The P(r) function was calculated using the Legendre method and the D_max_ value was determined through the likelihood search in Scatter.

---

**Fig. S1.**
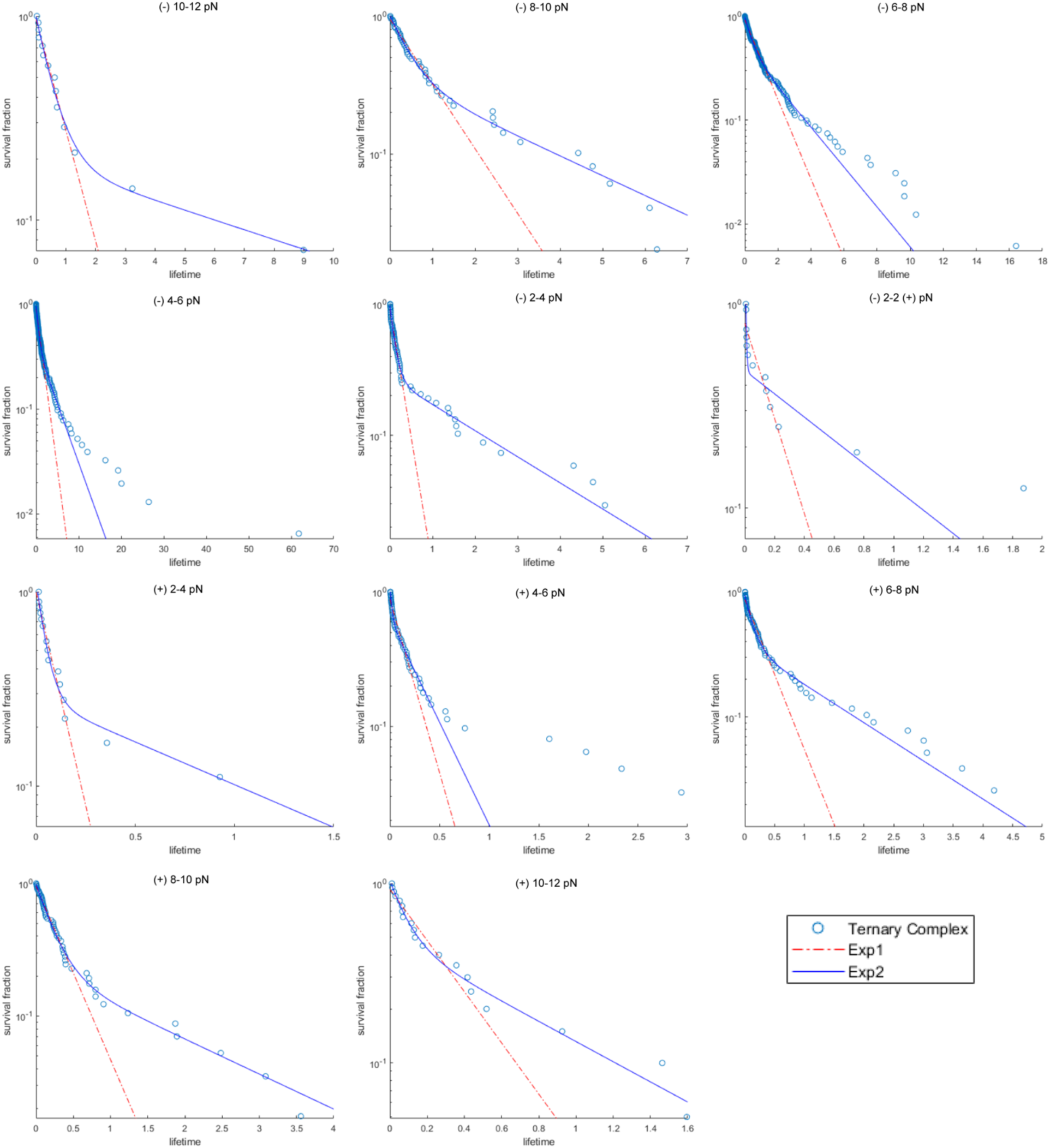
Survival plots of bound lifetimes for ternary complex. Each graph shows shows the survival fraction vs binding lifetimes in a bin of width 2 pN centered at the indicated load

**Fig. S2.**
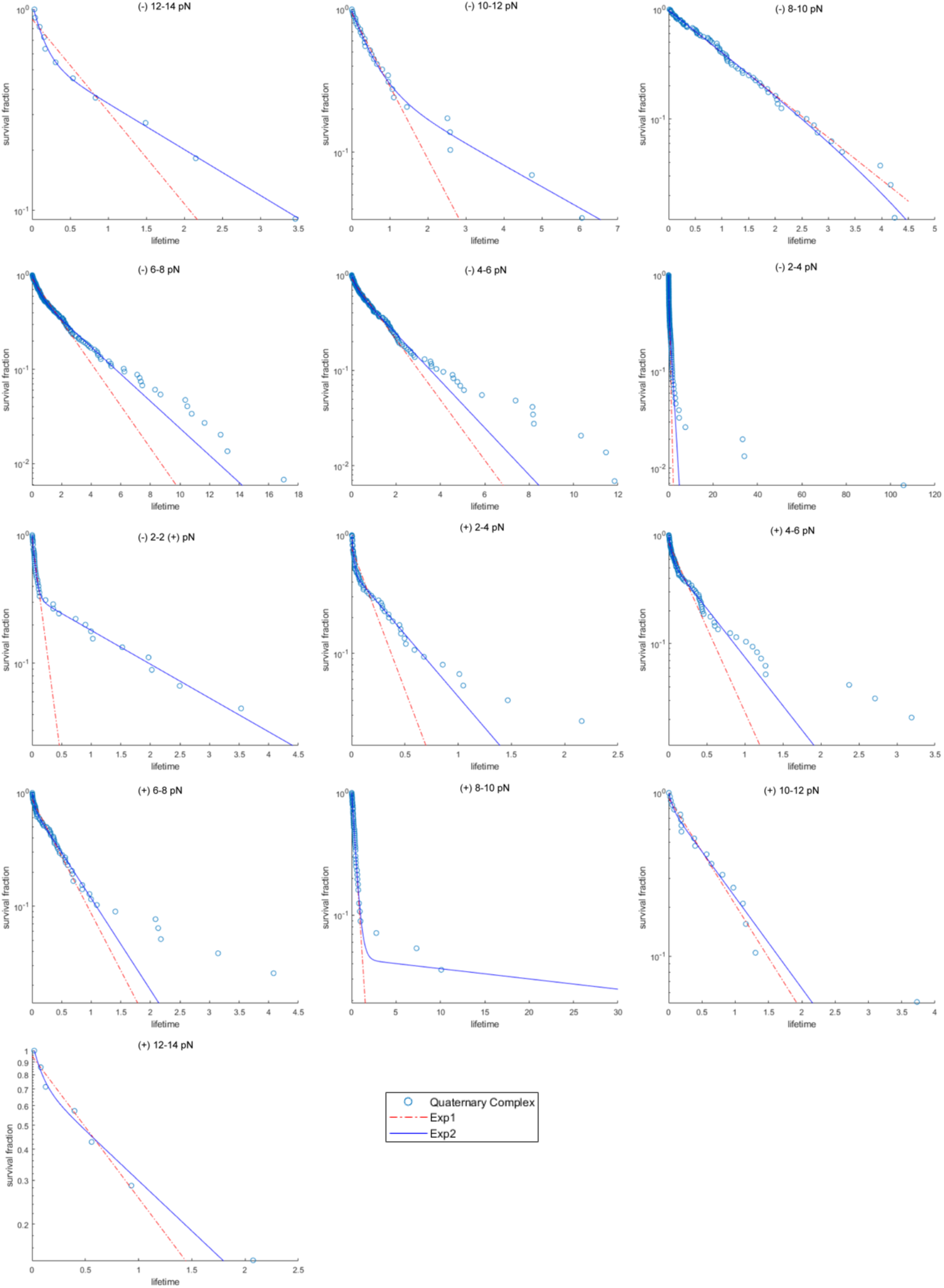
Survival plots of bound lifetimes for quaternary complex. Each graph shows the survival fraction vs binding lifetimes in a bin of width 2 pN centered at the indicated load.

**Fig. S3.**
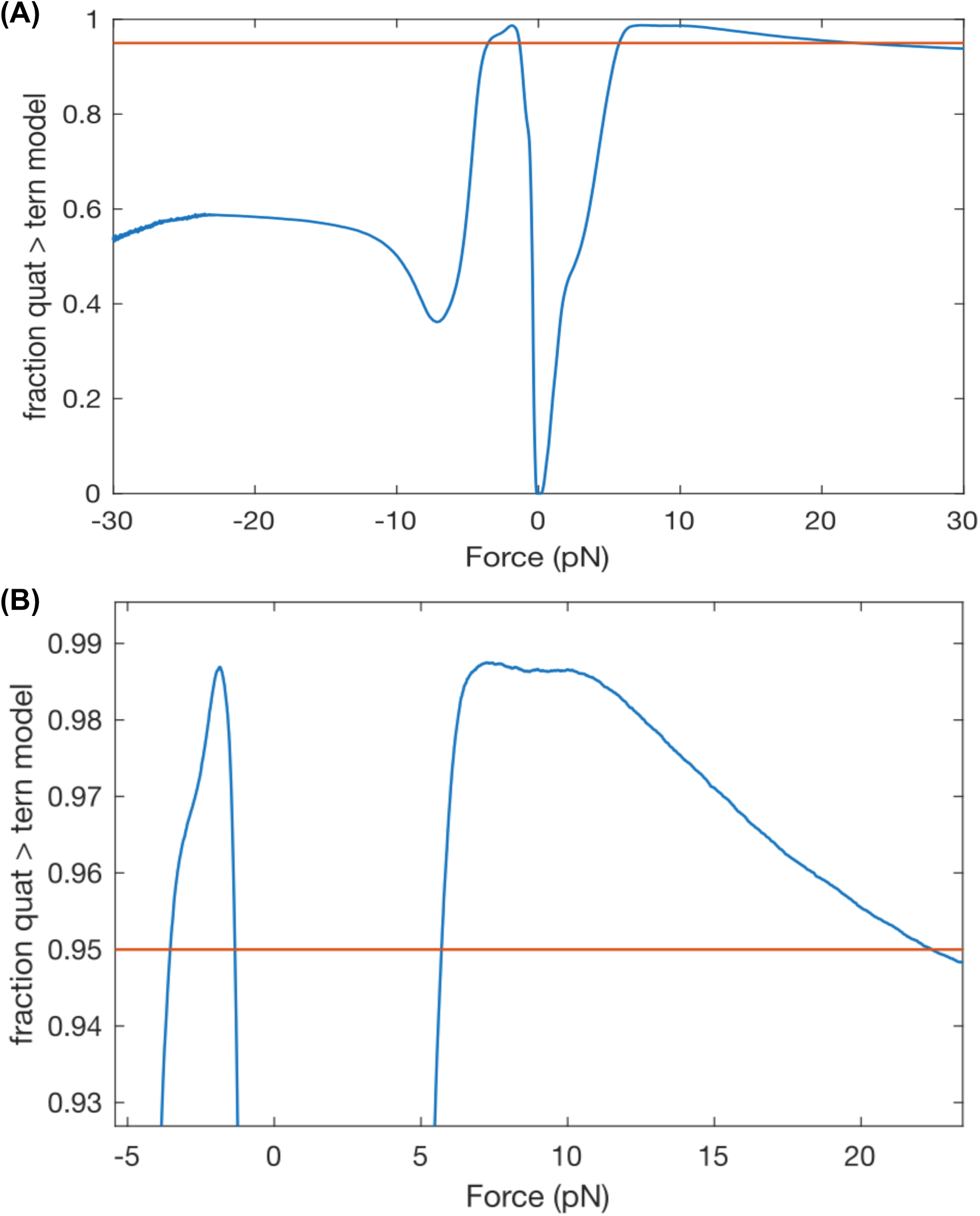
Model comparisons from bootstrapped parameters for the ternary and quaternary complexes. (A) Differences in lifetimes for 10^6^ randomly selected bootstrapped pairs quaternary and ternary models at forces between -30 and 30 pN. Regions where this fraction exceeds 95% suggests statistically meaningful differences in the models. (B) Zoomed-in view of the same graph as in (A).

**Fig. S4.**
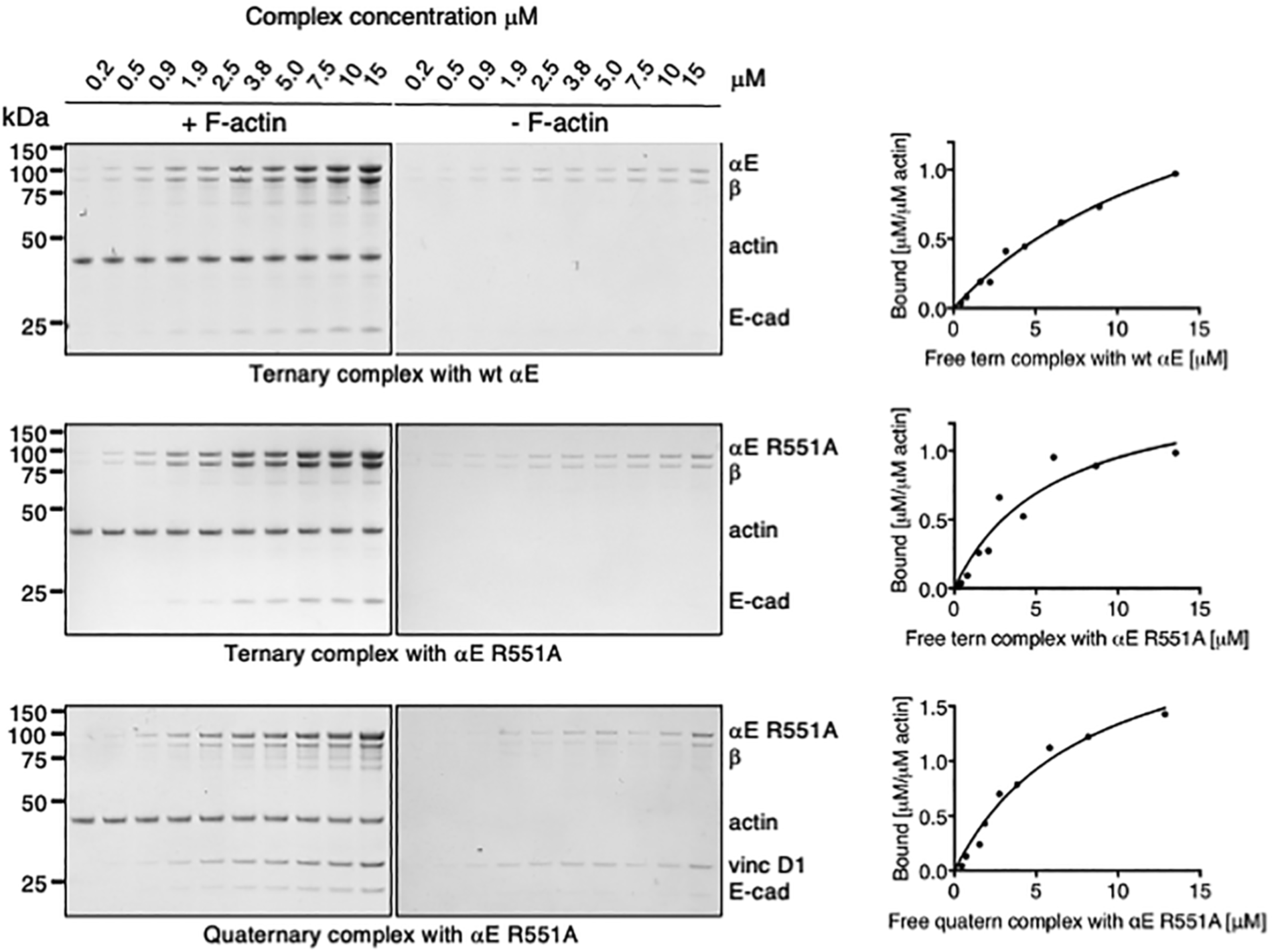
Actin binding of the ternary αE-catenin/ β-catenin/E-cadherin_cyto_ complex with wild-type αE-catenin and αE-catenin R551A and the quaternary αE-catenin R551A/β-catenin/E-cadherin_cyto_/vinculin D1 complex. SDS PAGE of actin pelleting assays in the presence and absence of F-actin are shown. Bound complex per actin is plotted against free complex concentration; a binding curve assuming a single binding site was determined using Graphpad. Two to four replicates were performed for each complex and a representative SDS gel with the corresponding binding curve is shown.

**Fig. S5.**
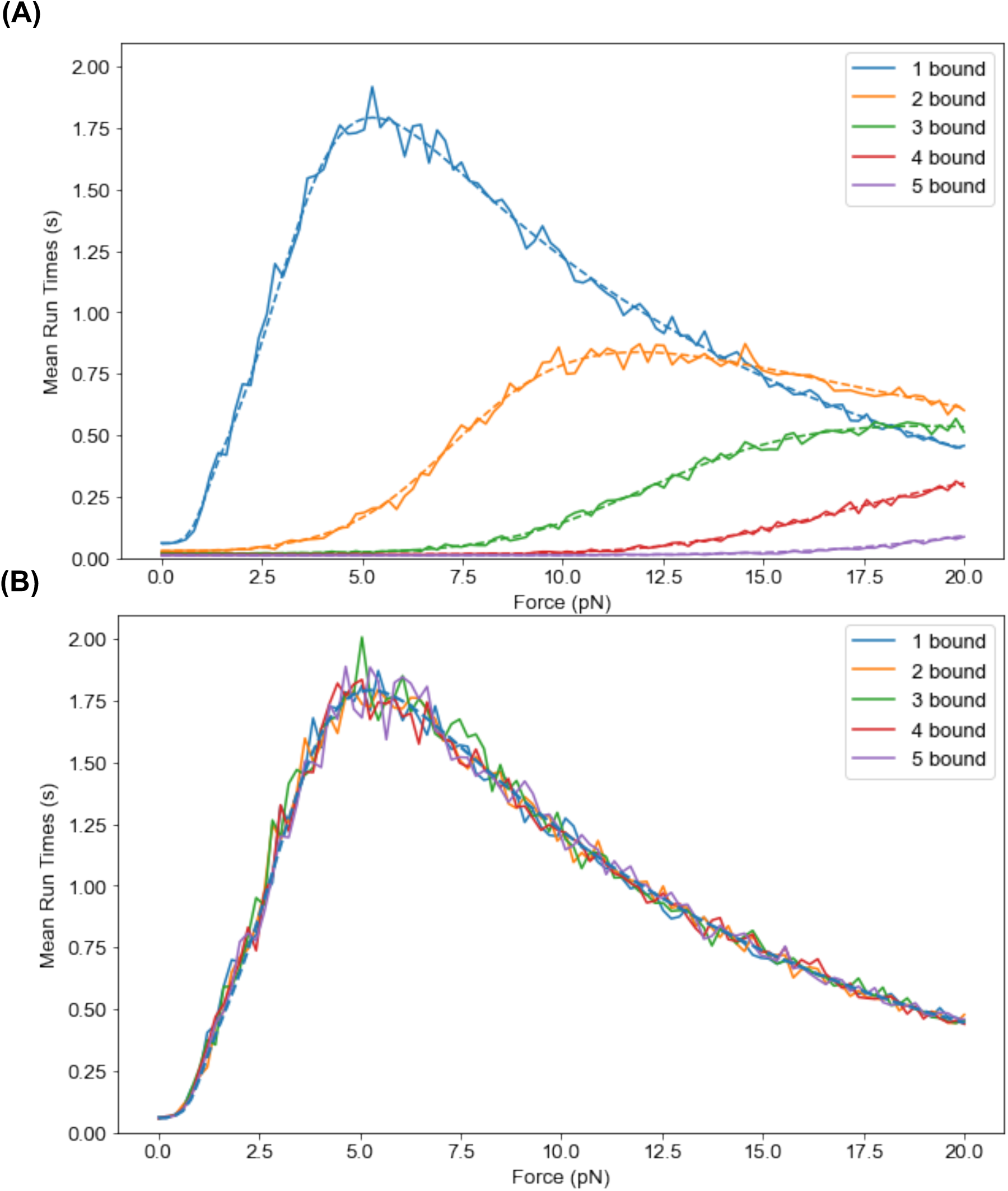
Analytical and simulated *N^th^* step mean bond survival time for quaternary^(-)^ complex. (A) Even load sharing. Monte Carlo simulations (—) are consistent with analytical (- -) treatment of bond survival times where bound complexes share equal load. The assumption that complexes share load equally result in a prediction where 〈*τ*〉*_N_* decreases as more complexes are bound to F-actin, which is inconsistent with experimental observations. (B) Unequal load sharing. Monte Carlo simulations (—) are consistent with analytical (- -) treatment of when only one complex is bound to F-actin. The assumption that one complex bears all the load predicts bond survival times that are independent of the number of interacting complexes.

**Fig. S6.**
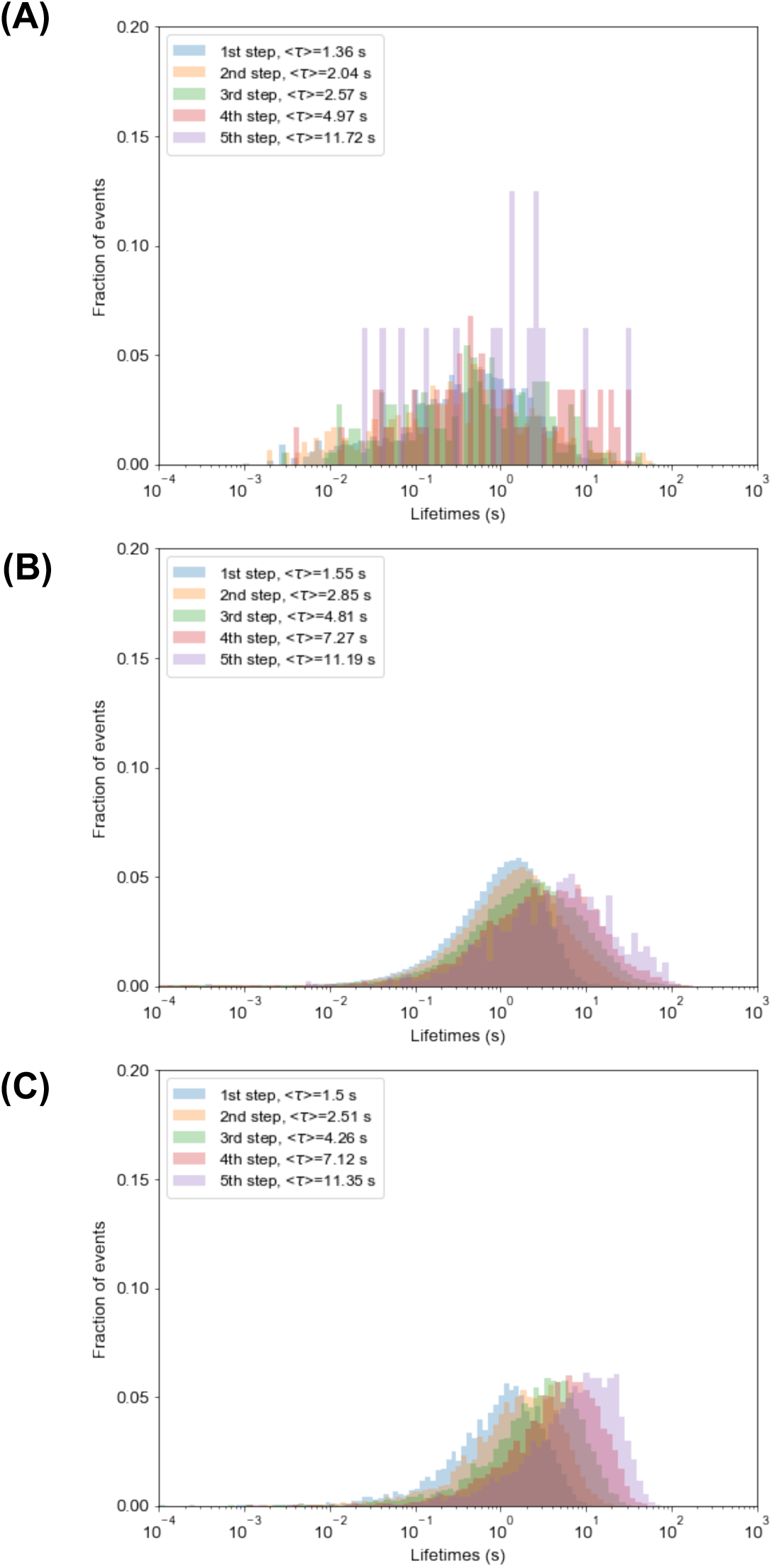
Distribution of average bound lifetimes in the quaternary^(-)^ complex. (A) Average bound lifetime distribution as stochastically sampled in OT experiments. Predicted average bound lifetime distributions for (B) neighbor-neighbor stabilization model for *p_o_* = 0.15, *k_on_* = 100 s^-1^, ΔE = 1.5 k_B_T, *F* = 8 pN and (C) geometric model for *Δx_20_* = -0.27.

**Fig. S7.**
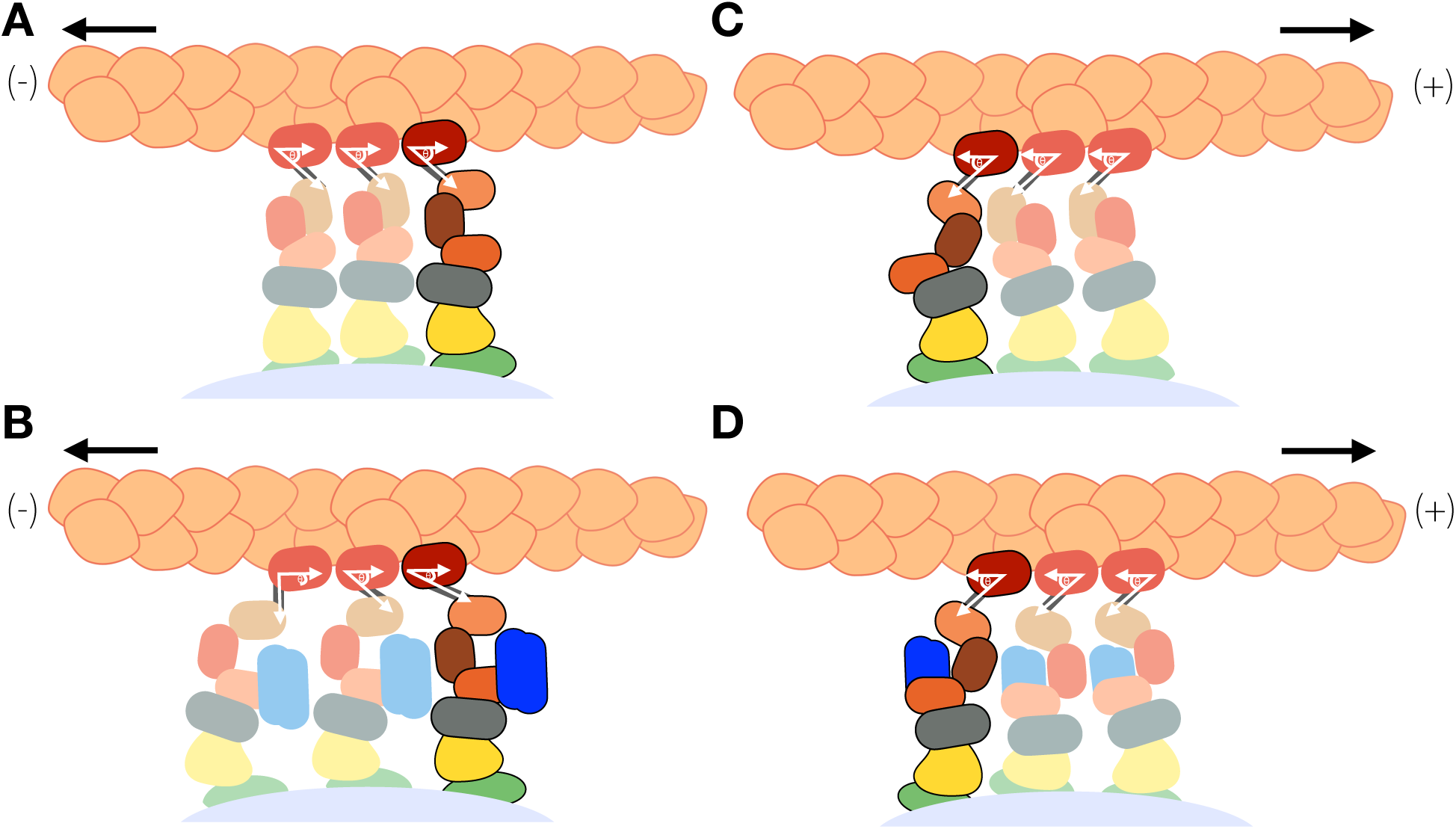
Geometric model as a possible mechanism for increased bound-state lifetimes for cadherin/catenin complexes with (-)-end directed force when vinculin is bound. The distance parameter *x_ij_* for a given transition in the two-state catch bond model depends on the projection of the force *F* onto the reaction coordinate: *x_ij_* = ‖*F*‖*r_ij_* ‖cos *θ*. Simultaneous binding by neighboring quaternary complexes may exert a torque on the F-actin filament, thus rotating *F* relative to *r_ij_* for the load-bearing complex. In simulations, of all the elementary rate constants in the two-state catch bond model, modest alterations in *x_20_^(-)^*, corresponding to detachment of the strongly bound state with load in the (-) direction, could result in appreciable changes in the bound-state lifetime: a 5° rotation in the axis of force application per bound complex would be sufficient to account for the increased lifetime.

**Fig. S8.**
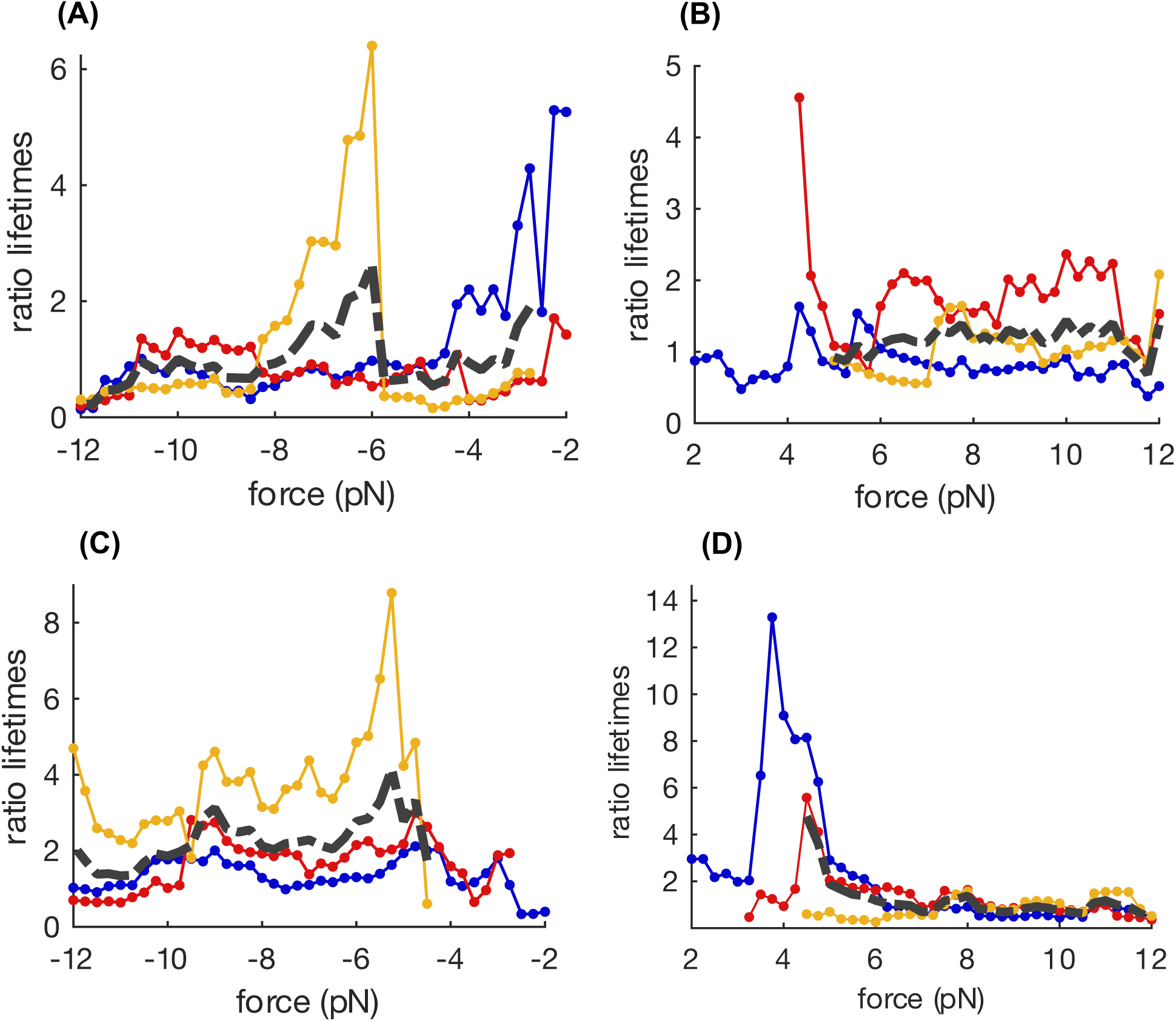
Ratio of *N^th^* step to last step average binding lifetimes vs force. (A) Ternary^(-)^ (B) Ternary^(+)^ (C) Quaternary^(-)^ (D) Quaternary^(+)^ complexes. Blue: 2^nd^-from-last vs last step, red: 3^rd^-from-last vs last step, yellow: 4^th^-from-last to vs last step, grey: overall average. For quaternary^(-)^, variations in mean binding lifetimes of the N^th^ step do not appear to vary as a function of load. Variability in 4^th^-from-last to vs last step likely reflects relatively low numbers of observations.

**Fig. S9.**
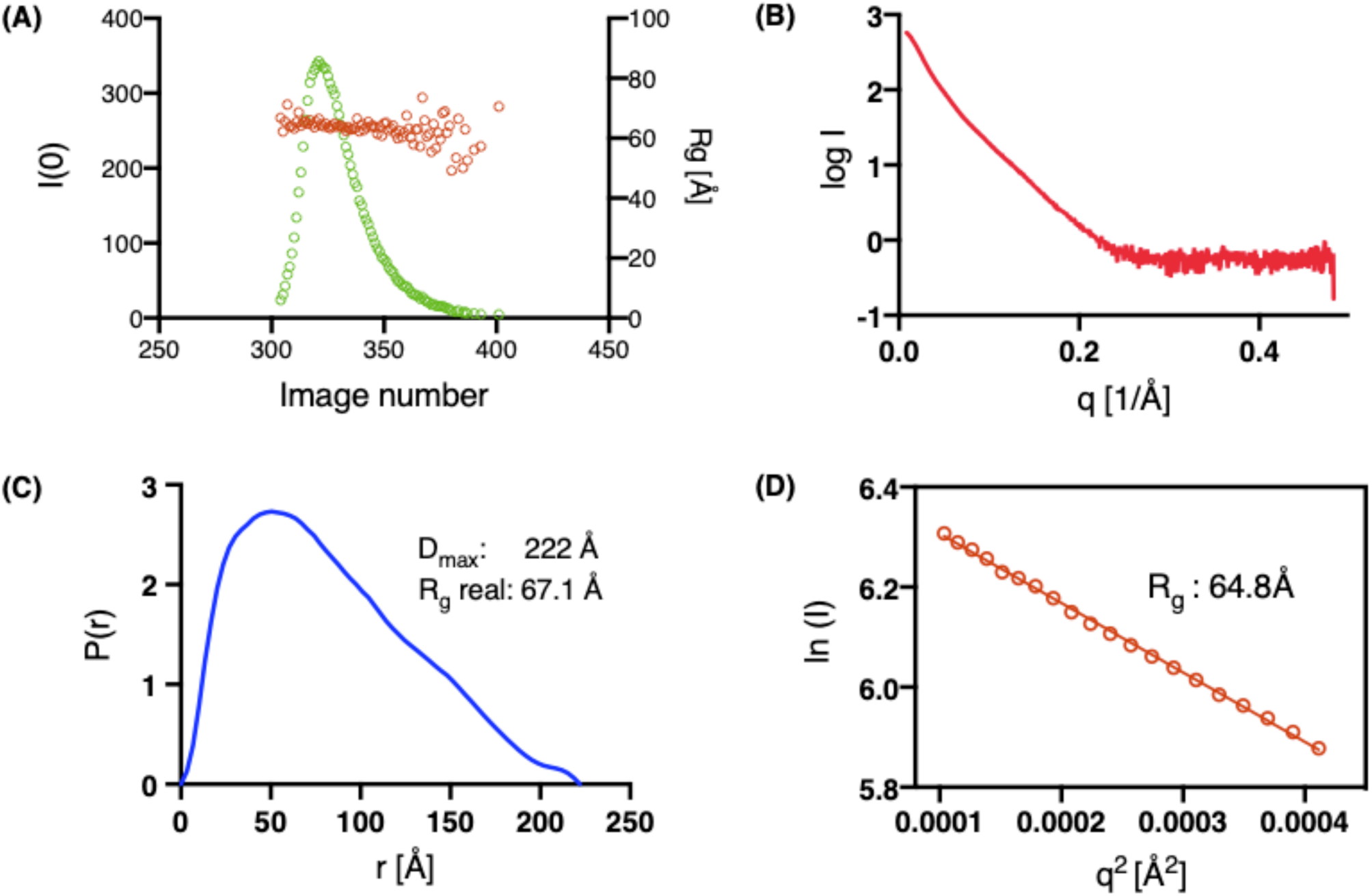
SAXS analysis of the ternary αE-catenin/μ-catenin/E-cadherin complex. (A) SEC-SAXS elution profile of the ternary complex with the extrapolated I(0) value plotted in green and the R_g_ value plotted in red. (B) Scattering curve of the ternary complex and (D) Guinier plot with the calculated R_g_ value. (C) Distance distribution function of the ternary complex with R_g_ and D_max_ estimates.

**Fig. S10.**
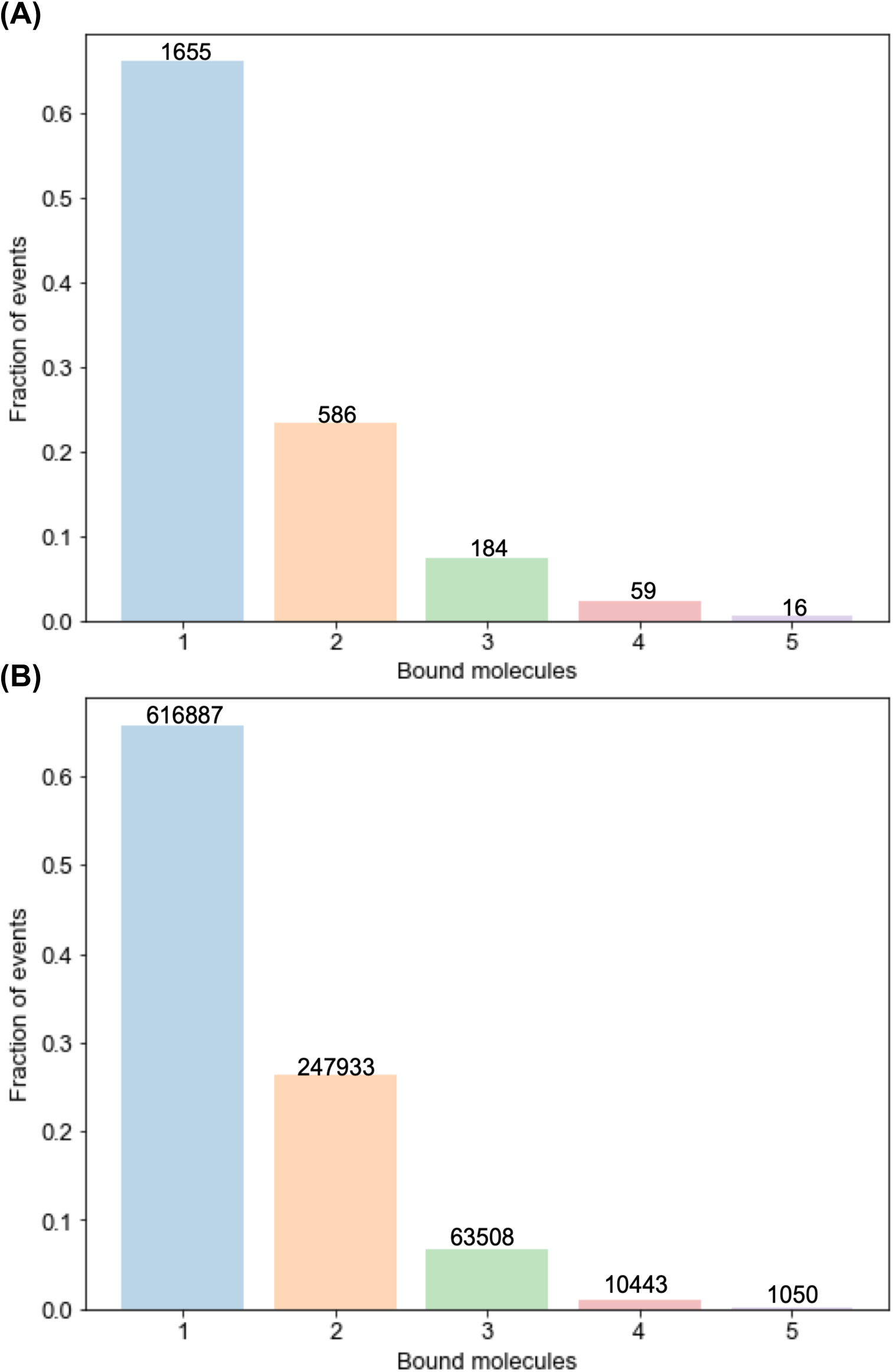
Power law distribution of N^th^ bound steps in the quaternary^(-)^ complex. (A) Experimentally determined N^th^ step distribution for the quaternary^(-)^ dataset. (B) N^th^ step event distribution in neighbor-stabilization models, which assumes 15% of complexes are active with a *k_on_* of 100 s^-1^ and ΔE of 1.5 k_B_T.

**Fig. S11.**
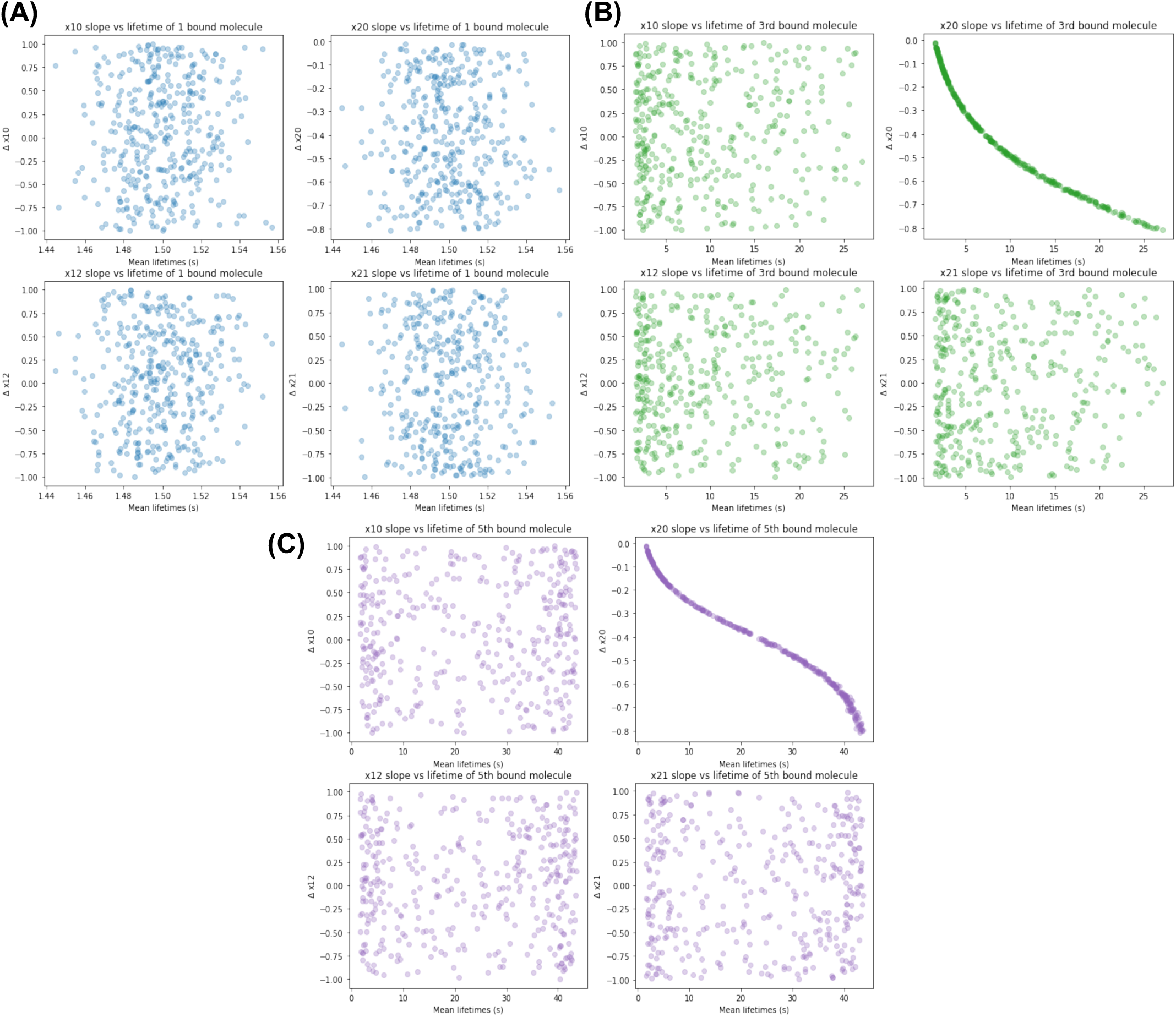
*Δx_ij_* sensitivity analysis for quaternary^(-)^ complex lifetimes vs *N*. (A) Binding lifetimes for one complex do not vary with distance parameters because the alignment angle *θ* is consistent with values derived from MLE fits. All mean lifetimes from simulations range from 1.44-1.56 s. (B) Simulations initialized with 3 molecules show that predicted binding lifetime increases as *x_20_* decreases. (C) Simulations initialized with 5 molecules show similar results reported in panel (B). The binding lifetimes increases as more molecules are initialized.

**Table S1.** Kinetic parameters for the two bound-state catch bond model for ternary and quaternary complexes.

**Ternary Complex**
| | $k_{ij}^0$ | CI ($s^{-1}$) | $x_{ij(-)}$ | CI (nm) | $x_{ij(+)}$ | CI (nm) |
| --- | --- | --- | --- | --- | --- | --- |
| **1 → 0** | 13.57 | (13.27, 14.21) | 0 | fixed | 0 | fixed |
| **2 → 0** | 0.22 | (0.15, 0.35) | 0.55 | (0.28, 0.78) | 0.98 | (0.74, 1.19) |
| **1 → 2** | 0.15 | (0.05, 0.39) | 4.72 | (3.92, 5.70) | 2.73 | (2.11, 3.40) |
| **2 → 1** | 6.27 | (1.61, 369.66) | 3.46 | (1.30, 18.30) | 15.00 | fixed |

**Quaternary Complex**
| | $k_{ij}^0$ | CI ($s^{-1}$) | $x_{ij(-)}$ | CI (nm) | $x_{ij(+)}$ | CI (nm) |
| --- | --- | --- | --- | --- | --- | --- |
| **1 → 0** | 17.50 | (17.26, 19.12) | 0 | fixed | 0 | fixed |
| **2 → 0** | 0.29 | (0.20, 0.36) | 0.42 | (0.30, 0.62) | 0.57 | (0.46, 0.72) |
| **1 → 2** | 0.51 | (0.19, 0.85) | 4.54 | (3.96, 5.56) | 2.49 | (2.12, 3.43) |
| **2 → 1** | 19.82 | (2.21, 40.29) | 15.00 | fixed | 9.82 | (1.01, 16.51) |

**Table S2.** Average lifetimes across all forces for 1^st^-5^th^ steps of quaternary and ternary complexes Ternary complex.

| **Ternary complex** |  |  |  |  |  |  |
| --- | --- | --- | --- | --- | --- | --- |
| Step Number | Negative |  |  | Positive |  |  |
|  | N | Avg F (pN) | Avg Dwell (s) | N | Avg F (pN) | Avg Dwell (s) |
| 1 | 455 | -5.98 | 1.59 | 245 | 6.82 | 0.44 |
| 2 | 455 | -7.86 | 1.19 | 245 | 9.20 | 0.33 |
| 3 | 149 | -9.12 | 1.12 | 74 | 10.45 | 0.61 |
| 4 | 54 | -10.09 | 1.09 | 38 | 11.73 | 0.33 |
| 5 | 14 | -11.38 | 0.46 | 18 | 12.34 | 0.61 |

| **Quaternary complex** |  |  |  |  |  |  |
| --- | --- | --- | --- | --- | --- | --- |
| Step Number | Negative |  |  | Positive |  |  |
|  | N | Avg F (pN) | Avg Dwell (s) | N | Avg F (pN) | Avg Dwell (s) |
| 1 | 586 | -5.88 | 1.58 | 359 | 5.99 | 0.66 |
| 2 | 586 | -8.07 | 2.04 | 359 | 8.51 | 0.91 |
| 3 | 184 | -8.64 | 2.57 | 104 | 9.10 | 0.93 |
| 4 | 59 | -9.03 | 4.97 | 38 | 9.70 | 0.69 |
| 5 | 16 | -9.99 | 11.72 | 11 | 10.89 | 0.93 |

**Table S3.** Mean binding lifetimes at low force.

|  | **N** | **Experimental**<br>**lifetime (ms)** | **Fast**<br>**rate (s<sup>-1</sup>)** | **Intermediate**<br>**rate (s<sup>-1</sup>)** | **Slow**<br>**rate (s<sup>-1</sup>)** | **Adjusted**<br>**lifetime (ms)** |
| --- | --- | --- | --- | --- | --- | --- |
| control | 35 | 10 | 180 | 40 | N/A | N/A |
| ternary | 90 | 62 | N/A | 30 | 2.5 | 76 |
| quaternary | 210 | 44 | N/A | 37 | 1.8 | 60 |

**Table S4.** 2D KS test with Bonferroni corrections comparing ternary vs. quaternary complexes for a given step number in each loading direction.

| Direction | 1<sup>st</sup> step | 2<sup>nd</sup> step | 3<sup>rd</sup> step | 4<sup>th</sup> step | 5<sup>th</sup> step |
| --- | --- | --- | --- | --- | --- |
| (-) | $1.62 \times 10^{-3}$ | $1.07 \times 10^{-9}$ | $1.14 \times 10^{-5}$ | $2.27 \times 10^{-3}$ | *$3.43 \times 10^{-2}$ |
| (+) | $2.55 \times 10^{-5}$ | $3.36 \times 10^{-7}$ | $5.78 \times 10^{-3}$ | $1.61 \times 10^{-3}$ | *$1.18 \times 10^{-1}$ |

\*Significance threshold for Bonferroni correction is 0.01

**Table S5.** Weighted bootstrap fitting of an exponential function for binding lifetime vs step number

|  | **c** | **95% CI** | **Adjusted R<sup>2</sup>** |
| --- | --- | --- | --- |
| ternary<sup>(-)</sup> | -0.24 | (-0.34, -0.14) | 0.90 |
| ternary<sup>(+)</sup> | -0.02 | (-0.19, 0.14) | 0.00 |
| quaternary<sup>(-)</sup> | 0.32 | (0.21, 0.44) | 0.86 |
| quaternary<sup>(+)</sup> | 0.06 | (-0.05, 0.16) | -0.16 |

Weighted fits to $L(n) = L_1 \exp(c \cdot (n - 1))$ , where $L(n)$ is the expected lifetime for a step number $n$ , where $L_1$ is the lifetime of one bound complex. $c$ is a parameter determined from weighted bootstrap fitting. Values for $c$ and $R^2$ for the ternary<sup>(+)</sup> and quaternary<sup>(+)</sup> datasets indicate that fits are indistinguishable from a flat line.

